# Plant recognition by *Trichoderma harzianum* elicits upregulation of a novel secondary metabolite cluster required for colonization

**DOI:** 10.1101/2023.04.12.536597

**Authors:** Miriam Schalamun, Guofen Li, Wolfgang Hinterdobler, Dominik K. Großkinsky, Stéphane Compant, Assia Dreux-Zigha, Jennifer Gerke, Russell Cox, Monika Schmoll

**Affiliations:** AIT Austrian Institute of Technology GmbH, Center for Health and Bioresources, Konrad Lorenz Strasse 24, 3430 Tulln, Austria; Greencell, 63360 St Beauzire, France; Leibnitz University Hannover, Institute of Organic Chemistry, Schneiderweg 38, 30167 Hannover, Germany; University of Vienna, Department of Microbiology and Ecosystem Science, Division of Terrestrial Ecosystem Research, Djerassiplatz 1, 1030 Vienna, Austria

**Keywords:** *Trichoderma harzianum*, *Hypocrea lixii*, biocontrol, plant protection, plant-fungus interaction, secondary metabolism, MAMP (microbe-associated molecular pattern), interkingdom communication, PCA cluster

## Abstract

*Trichoderma harzianum* is a filamentous ascomycete frequently applied as biocontrol agent in agriculture. While mycoparasitism and antagonism of *Trichoderma* spp. against fungal pathogens are well known, early fungal responses to the presence of a plant await broader investigation. Analyzing early stages of plant-fungus communication we show that *T. harzianum* B97 chemotropically responds to a plant extract and that both plant and fungus alter secondary metabolite secretion upon recognition. We developed a strategy for omics-analysis simulating conditions of early plant recognition eliciting a chemotropic response in the fungus and found 102 genes to be differentially regulated, including nitrate and nitrite reductases. Additionally, the previously uncharacterized *<u>P</u>*lant *<u>C</u>*ommunication *<u>A</u>*ssociated (PCA) gene cluster was strongly induced upon recognition of the plant, comprises a palindromic DNA motif and was essential for plant colonization. The PCA-cluster is only present in the Harzianum clade of *Trichoderma* and closely related to a homologous cluster in *Metarhizium* spp. Horizontal gene transfer (HGT) was detected for PCA-cluster genes by plants, while the cluster in *T. harzianum* is likely under balancing or positive selection.

Hence, the PCA-cluster mediates early fungus-plant chemical communication and may be responsible for the high potential of *T. harzianum* and closely related species for biocontrol applications.

**Plain language summary:** Interactions of plants with fungi – beneficial or pathogenic – are crucial for the ecological function of both partners. Yet, the chemical “language” they use and how or when they use it is still insufficiently known. We describe discovery of a novel gene cluster, which is strongly induced upon plant recognition and essential for plant-fungal interkingdom interaction in the biocontrol-agent *Trichoderma harzianum*.

## Introduction

Natural environments harbor a complex community of microorganisms, which fulfill crucial tasks in the carbon cycle and can interact with plants as symbionts [1,2] or pathogens [3]. Climate change and global warming are bringing increased disease pressure, abiotic stresses and promote invasion of plant pathogens in new habitats [4,5]. Hence, better understanding for knowledge based application of biocontrol agents and biostimulants is required [6,7]. Fungi evolved elaborated mechanisms for dealing with their biotic and abiotic environment in terms of sensing and signaling mechanisms as well as strategies for effective competition and antagonism [8,9]. Fungal secondary metabolites are thereby of crucial importance for interactions with other microbes, animals and also with plants [10]. The abilities of some fungi to antagonize and kill their competitors is applied for protection of plants against pathogens and fungi of the genus *Trichoderma* are among the most broadly applied for this purpose [11]. Consequently, this genus also dominates research towards mycoparasitism, plant protection and biocontrol of plant pathogens [12,13].

Fungi of the genus *Trichoderma* [14,15] are typical inhabitants of the rhizosphere and are found in soils worldwide [16]. Several *Trichoderma* species are known as efficient biocontrol organisms and act as important symbionts with plants [17,18]. They are studied in detail for their capabilities in producing antibiotics, parasitizing other fungi – predominantly plant pathogens – and to compete with deleterious plant pathogens [13,16]. Beneficial *Trichoderma* strains further induce root branching, can increase shoot biomass and trigger systemic resistance as well as plant nutrient uptake [19].

Among the most important functions for successful fungal plant interaction – beneficial or pathogenic -is the ability to colonize plant roots [13,20]. *Trichoderma* spp. are able to efficiently colonize plant roots, although they mostly remain at the outer layers of the plant tissue [13,19]. Nevertheless, also truly endophytic strains of *Trichoderma* [21,22] are known to associate with plants, including also some *T. harzianum* strains [21,23]. Fungi communicate with their environment using a broad array of signals [24–26]. Chemical communication between fungi and plants is essential for interaction and diverse secondary metabolites are known to play an important role in this interplay [26]. Intriguingly, stressed plants were found to secrete specific compounds attracting beneficial fungi [27]. Overall, a considerable number of effectors [12] and secondary metabolites [19] including volatile organic compounds [28] are already known to contribute to successful biocontrol. In the plant, recognition of beneficial fungi like *Trichoderma* spp. leads to metabolic changes [29] and the onset of systemic resistance [30]. Acquisition of nutrients is supported by plant-fungus interactions, also if the requirements for mycorrhiza are not fulfilled.

One major question of the last decades was how fungi sense the presence of a plant. A seminal study on plant recognition by the fungal pathogen *Fusarium oxysporum* provided groundbreaking insights in this respect [31,32]. It was shown that this fungus chemotropically responds to the presence of a plant and that this response is dependent on a pheromone receptor of the fungus, which obviously senses a peroxidase of the plant [32,33]. This research and the developed method open up new possibilities to study plant-fungus as well as other intra-and interkingdom interactions and their determinants [34].

Biocontrol of plant pathogens [35] is a complex mechanism involving processes from secretion of enzymes to production of secondary metabolites to mycoparasitism on the fungal pathogen [36]. *Trichoderma* spp. as well as other fungi applied in agriculture as plant beneficial agents produce a broad array of secondary metabolites [37,38]. Nevertheless, these fungi (except for a few more problematic species like *T. brevicompactum*) have a long history of safe application worldwide. No contamination of treated crops has been observed and hardly any negative effects on plants are known. Thereby, not only the fungi themselves, but also their secondary metabolites can be applied in agriculture [39]. Moreover, investigation of secondary metabolites of *Trichoderma* and their functions bears the opportunity to identify novel, bioactive compounds potentially useful in medicine and industry [40]. Interestingly, evolutionary analysis revealed that the core genome of *Trichoderma* species comprises about 7000 genes and that a considerable number of genes crucial to their well-known functions in litter degradation or secondary metabolism was acquired by horizontal (or lateral) gene transfer (HGT) [14,41,42]. This successful sourcing of advantageous genes by *Trichoderma* is assumed to be corroborated by their capability of mycoparasitism, which brings them in contact with foreign DNA [41]. Such feeding activities along with physical association over prolonged periods of time, like for example an association of a fungus with a plant are known to facilitate HGT between eukaryotes [43]. Additionally, asexual development of fungi, involving unicellular spores, is in agreement with the “weak link hypothesis” for entry of foreign genes into eukaryotic genomes [44]. Acquisition of secondary metabolite clusters can be particularly beneficial for a biocontrol agent, due to their potential function in communication with the plant and/or fending of competitors or antagonizing pathogens. Nevertheless, the relevance of HGT in eukaryotes remains controversial [45,46], and careful analysis is required to exclude for example sequencing artifacts jeopardizing the results [47].

*Trichoderma harzianum* B97 [48] was selected for its high efficiency in stimulation of plant growth. Moreover, the strain shows solubilization of phosphate and can alleviate abiotic stresses. Due to its proven efficiency in agricultural applications, it was the ideal isolate for studying its communication with a plant in more detail. We show that *T. harzianum* B97 chemotropically reacts to the presence of a plant and chemically communicates with a living plant as well. Moreover, we found that the ability of B97 to efficiently colonize plant roots depends on a secondary metabolite cluster specifically induced during early plant recognition, which is likely under balancing or positive selection.

## Results

### T. harzianum is an efficient colonizer of plant roots

*T. harzianum* B97 was isolated from agricultural soil in France [48] and selected for its plant beneficial characteristics. Good colonization of plants by the fungus would indicate efficient communication with the plant [49]. Therefore, we studied, whether *T. harzianum* B97 would be able to efficiently colonize plant roots. We analyzed colonization after co-inoculation of wheat seedlings with *T. harzianum* B97. Staining of roots with wheat germ agglutinin (WGA)-AlexaFluor488 and confocal microscopy showed efficient colonization of roots by *T. harzianum* B97 (Figure 1A).

**Figure 1.**
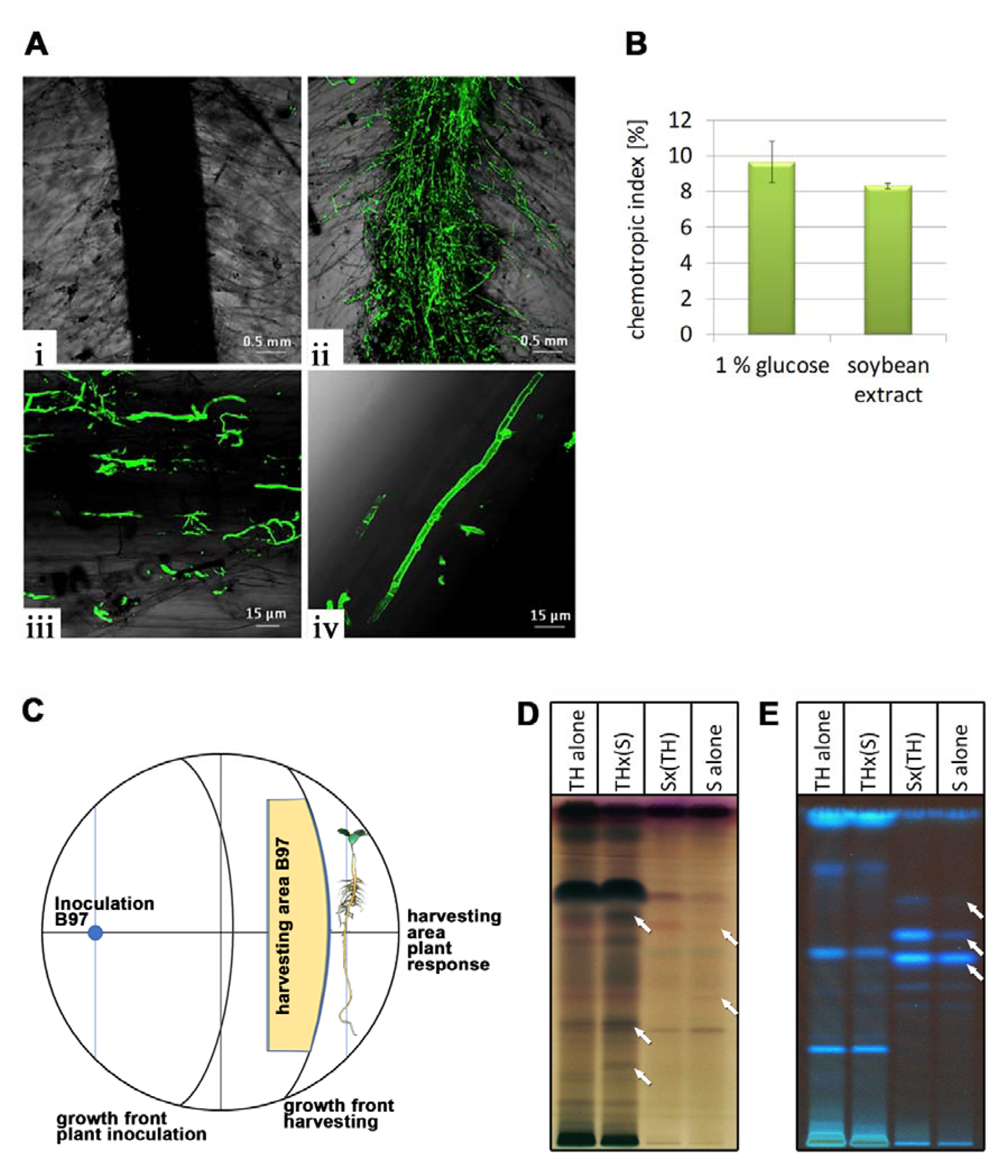
Assessment of *T. harzianum* B97 – plant interaction. (A) CSLM Microphotograph of uninoculated control roots of wheat (i) and wheat roots with *T. harzianum* B97 (ii-iv) at the root hair zone and stained with WGA-Alexa Fluor488® showing B97 as green fluorescent colonizing wheat root hairs (iii) or the root surface (iv). (B) Chemotropic indices of *T. harzianum* B97 to the presence of 1 % (w/v) glucose or root exudates of soy plant. Analyses were done in biological duplicates, at least 400 germlings were counted per experiment. (C) Schematic representation of the experimental setup for analysis of plant-fungus communication. Plants were allowed to interact with the fungus for 13 hours and harvesting was done before contact. Mycelia for investigation of the transcriptome was isolated from the mycelial growth front (“harvesting area B97”, yellow). For secondary metabolite analysis by HPTLC the agar slice including cellophane overlay from exactly the same area was excised. For analysis of the response of the plant an agar slice on the other side of the root was excised in order to avoid interference with fungal metabolites. For control plates the setup and positioning of harvesting areas was exactly the same. (D, E) HPTLC analysis of *T. harzianum* B97 alone on the plate (TH alone), *T. harzianum* B97 in the presence of the root of soy plant (THx(S)), the root of soy plant in the presence of the fungus (Sx(TH)) and the root of soy plant alone (S alone). Two different visualizations are provided (D: Visible light after anisaldehyde derivatization and E: Remission at 366 nm) and show differentially secreted metabolites between interaction partners alone and in combination. A representative analysis result is shown. For results of all three biological replicates, see supplementary file 2, figure S1.

### T. harzianum B97 shows chemotropic response to soybean root exudates

Recognition of the presence of the plant in the vicinity is crucial for initiation of interaction. Moreover, successful interaction with different plant species is a desirable trait for biocontrol agents. Recently, attraction of a *T. harzianum* strain to plant roots was shown [27]. Hence we wanted to test first, whether *T. harzianum* B97 chemotropically reacts to the presence of roots of soy plants using their root exudates as chemotropic agent. Optimization of the assay to *T. harzianum* B97 yielded an optimal working concentration of 0.0025 % peptone from casein to support germination without inducing multipolarity. We tested the response of *T. harzianum* B97 to 1 % (w/v) glucose, which resulted in a chemotropic index of 9.66 ±1.15 % and was hence in the range seen previously for fungi [32]. Subsequent analysis of the chemotropic response to soybean root extracts showed a chemotropic index of 8.32% ± 0.15%, representing low but clearly present response (Figure 1B). This results further shows that the previously detected chemotropic response is not limited to the *Fusarium*-tomato interaction system, but a rather general response.

### Secretion of secondary metabolites changes in the presence of soy bean roots

As *T. harzianum* B97 clearly reacts to the presence of the plant, we wanted to test whether chemical communication is initiated as a consequence of recognition. We used conditions as close as possible to those applied in the chemotropic assay, with a low level of carbon source (0.1 % (w/v) glucose) and minimal medium nutrients to support growth and still allow plant recognition (Figure 1C). After 34 hours of growth of *T. harzianum* B97, roots of soybean plants were placed in 3 cm distance of the fungal growth front and incubated for 13 hours in the darkness to enable communication. To ensure that communication occurs via the medium or via volatile organic compounds, but not due to direct contact, we only used plates where fungus and plants remained without physical contact at the time of harvesting. Thereafter, agar slices were excised from the area covered by the fungus for assessing changes in fungal metabolite profiles as well as from the area opposite of the plant root to analyze alterations in metabolites secreted by the plant (Figure 1C). Secondary metabolite patterns from the fungus grown without a plant and of the plant in the absence of the fungus under otherwise similar conditions were used as controls.

Indeed, after 13 hours of exposure of the fungus to the plant, we observed additional bands appearing in the high performance thin layer chromatography (HPTLC) analysis, reflecting a reaction of the fungus to the plant (Figure 1D,E). Also the plant secreted additional compounds upon detection of the fungus, which were not present in the assay without the fungus (Figure 1D,E). Consequently, *T. harzianum* B97 initiates chemical communication with the soy plant roots within 13 hours of co-cultivation.

### Transcriptome analysis of early stages of plant recognition

Having confirmed that plant recognition by *T. harzianum* B97 indeed occurs and elicits a two-way communication, we adapted transcriptome analysis to these conditions by covering the agar surface with cellophane to enable harvesting of the mycelium. Due to the high similarity of *T. harzianum* B97 with the previously sequenced reference strain *T. harzianum* CBS226.95 [41] we refer to protein IDs from the respective public genome database at JGI (https://mycocosm.jgi.doe.gov/Triha1/Triha1.home.html) in the following.

Since transcript levels reflect investment of resources for functions important under a certain condition, we checked which functions were represented under the conditions used for secondary metabolite screening and transcriptome analyses. Functional category analysis of the 250 genes with highest transcript levels under the applied conditions showed considerable investments in metabolic functions, energy production and transport among others (Figure 2A).

**Figure 2.**
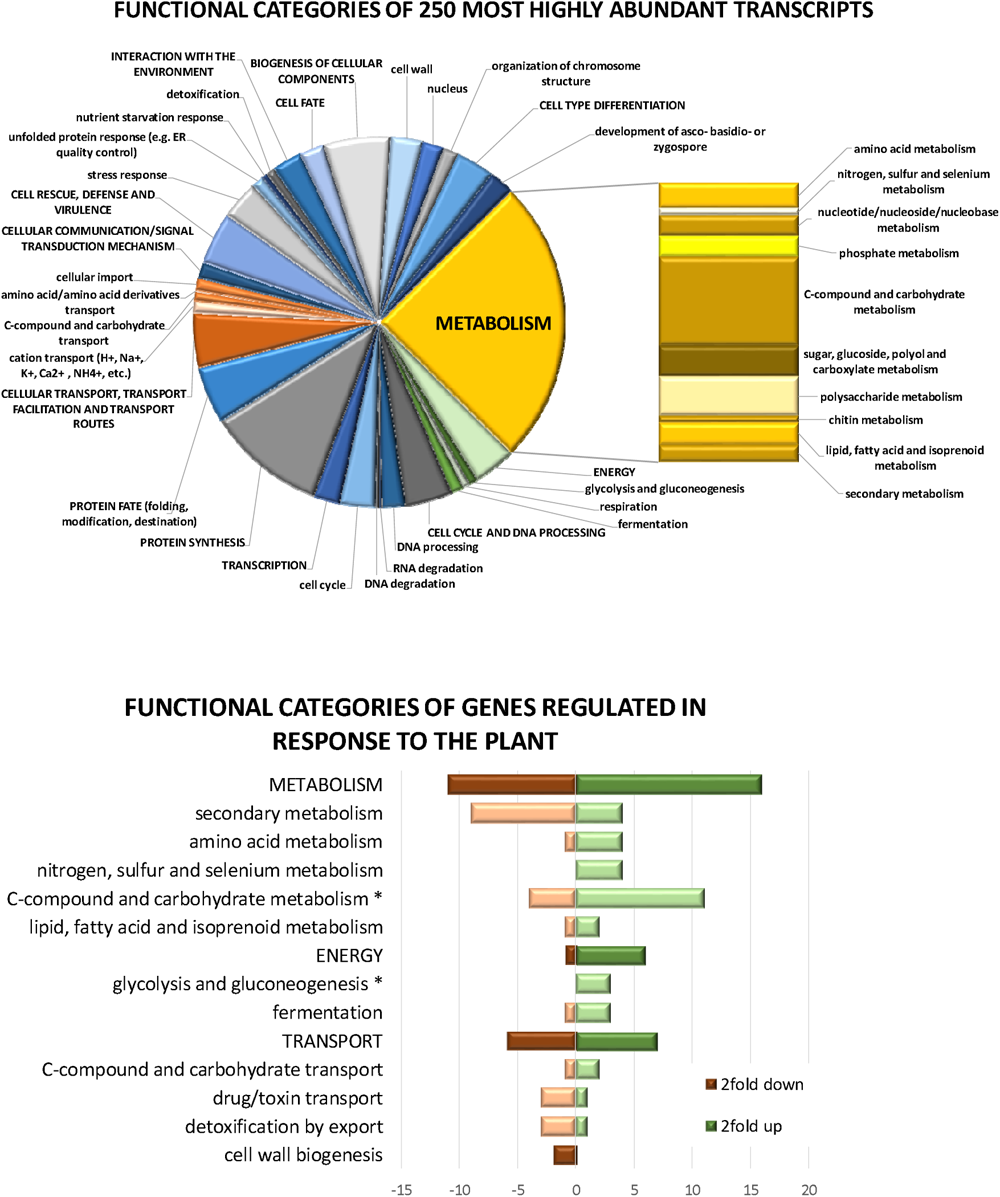
Functional analysis of gene expression in *T. harzianum* B97. (A) Functional categories represented among the 250 most abundant transcripts under the conditions of simulated chemotropic response. (B) Major functional categories assigned to genes differentially regulated in the presence of a soy plant. Significantly enriched categories are marked with an asterisk.

Besides the expected enrichment in metabolic functions, energy metabolism and carbohydrate metabolism, we also found that the highest levels of transcript abundance were enriched in functions in stress response (p-value 1.96e-03) and unfolded protein response (p-value 7.89e-04). Additionally, also polysaccharide metabolism was significantly enriched in this highly transcribed gene set (p-value 2.49e-11), with the homologues of the cellobiohydrolases *cbh1* and *cbh2* comprised in this group. These patterns reflect that the chosen condition indeed represent an environment of low nutrient availability inducing cellobiohydrolases likely due to starvation. Consequently, this condition closely resembles the conditions present in the chemotropic assay.

### Specific gene regulation in the presence of a plant

Comparison of genes differentially regulated between growth alone on the plate and in the presence of a plant confirmed that our experimental setup captured a very early specific stage of plant recognition by *T. harzianum* B97 and likely represents the onset of communication. In total, only 102 genes were significantly (p-value <0.01) regulated more than 2fold (41 down, 61 up) upon recognition of the plant (supplementary file 1), which share functions in energy production, metabolism and transport (Figure 2B). Genes upregulated in the presence of the plant were enriched in functions of C-compound and carbohydrate metabolism (p-value 1.04e-03), glycolysis and gluconeogenesis (p-value 4.09e-03) and electrochemical potential driven transport (p-value 4.00e-03). Interestingly, the gene set down-regulated in the presence of the plant is enriched in functions in secondary metabolism (p-value 5.03e-04) and drug/toxin transport (p-value 4.26e-03).

Specifically, we detected a nitrate reductase (Triha_507858) and a nitrite reductase gene (Triha_507859) to be up-regulated 17fold or 4fold, respectively upon plant recognition. These genes represent the homologues of the *Aspergillus nidulans* genes *niiA* and *niaD*, which share a bidirectional promotor and play important roles in nitrogen uptake and metabolism [50,51]. However, the putative homologue of *crnA*, the major facilitator superfamily (MFS) transporter gene associated with this cluster in *A. nidulans* (Triha1_142220) and the major transcription factor genes responsible for regulation of nitrogen metabolism, *areA* (Triha1_451) and *areB* (Triha1_70872), are not significantly differentially regulated under these conditions.

Considering fungus-plant interaction, also the more than 5fold up-regulation of Triha_398864, encoding a homologue of Epl1/Sm1 is interesting. These ceratoplatanin-like proteins play a role in colonization of plant roots and as effectors [52]. In *T. harzianum*, Epl1 regulates virulence of the plant pathogen *Botrytis cinerea*, mycoparasitism as well as plant immunity at early stages of root colonization [53,54], which is in perfect agreement with our hypotheses.

Furthermore, transcript abundance of two predicted protease genes (Triha1_541862; 5.3fold down and Triha1_98848; 2.2fold down) is decreased in the presence of plant roots. As two putative terpene synthase genes, *tps1* (Triha1_497584; 2.8fold up) and *tps11* (Triha1_523651; 3.1fold up) are up-regulated, a role of terpenoid compounds in plant interaction is worth further investigation. Additionally, an as yet uncharacterized non-ribosomal peptide synthase (NRPS, Triha_155805) and a putative polyketide synthase (PKS, Triha_546993) are more than 2 fold upregulated upon plant sensing.

A further, strongly upregulated gene (32fold) is Triha1_36398, which is still uncharacterized and its encoded protein comprises no known domains. However, analysis of putative protein interaction partners using the homologue of this protein in *T. reesei* using the STRING database ([55]; https://string-db.org; version 11.5) suggests a connection to a predicted ferric reductase, which fits to its genomic vicinity next to a putative ferric reductase (Triha1_76871) in *T. harzianum*, which is also strongly upregulated and fits to the general picture of gene regulation in B97 upon plant recognition.

We conclude that the recognition of a plant in the environment causes *T. harzianum* to modulate secondary metabolism, but to also elevate certain metabolic capabilities, notably also nitrogen metabolism which might be beneficial for nutrient exchange with a plant. Although functions in C-compound and carbohydrate metabolism are enriched among up-regulated genes, this gene set does not include the common plant cell wall degrading enzymes. The high expression level of cellulases detected in all analyzed samples (see above) is not significantly altered upon recognition of a plant.

### A secondary metabolite cluster strongly up-regulated upon plant recognition

Despite the low number of regulated genes, we still found a strongly regulated gene cluster (Figure 3A, B), which is silent when the fungus is growing alone, and strongly induced upon recognition of the plant with up to 1000-fold upregulation (Figure 3B; supplementary file 1). This gene cluster has a size of 15 kb and is located on scaffold 23: 272 000 – 287 000 in the reference strain *T. harzianum* CBS226.95 and comprises all seven genes present in this area. We termed the cluster *<u>P</u>*lant *<u>C</u>*ommunication *<u>A</u>*ssociated (PCA) cluster which is comprised of Triha1_323871/*pca1*, Triha1_513502/*pca2*, Triha1_513502/*pca3*, Triha1_99174/*pca4*, Triha1_513504/*pca5*, Triha1_513505/*pca6* and Triha1_513506/*pca7*. None of the genes in the cluster was previously characterized.

**Figure 3.**
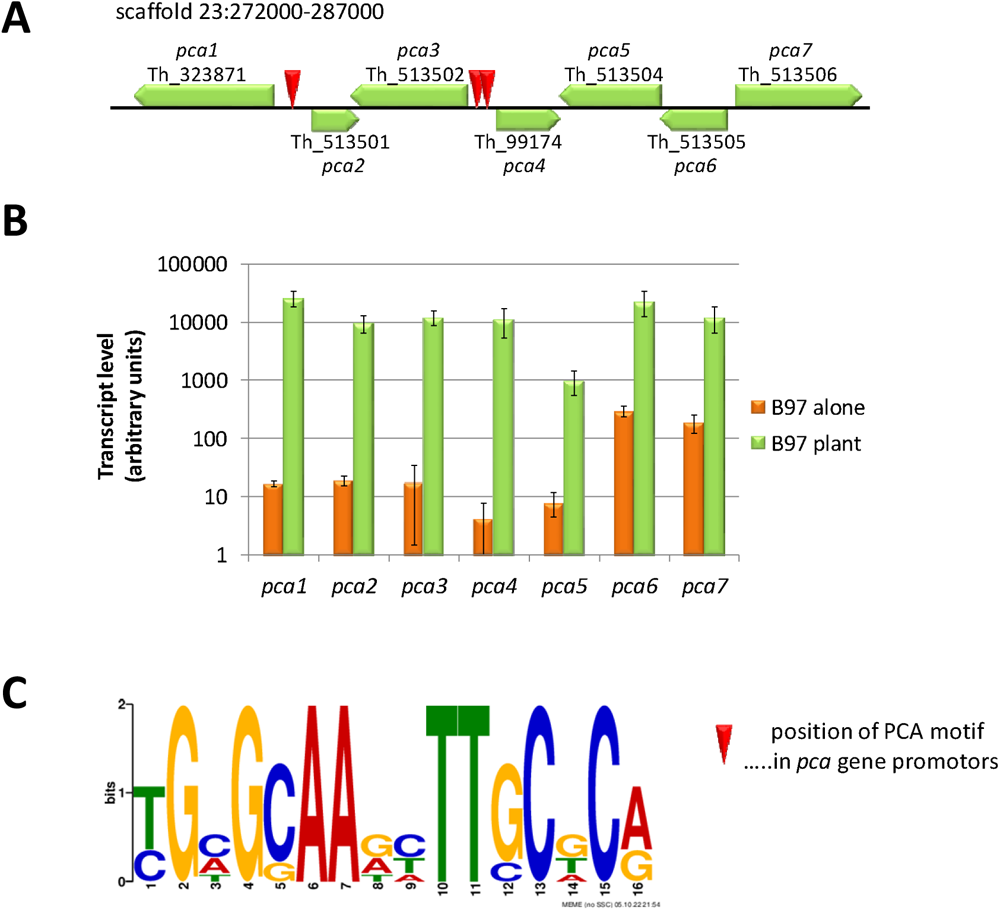
Schematic representation of the PCA cluster (A) and its regulation upon recognition of the plant (B). (A) Localization of *pca*-genes in the *T. harzianum* genome (JGI mycocosm; https://mycocosm.jgi.doe.gov/mycocosm/home). Approximate position of the PCA-DNA motif is shown with red triangles. (B) RPKM values of transcript levels of *pca* genes upon growth alone on the plate (orange bars), where transcripts were at very low basal levels (logarithmic scale is shown). Green bars represent transcript abundance upon recognition of the plant. Values represent means of three biological replicates and error bards show standard deviations. In all cases differential regulation is statistically significant (p-value <0.01) (C) PCA motif as found in the potentially bi-directional promotors of *pca1*/*pca2* and *pca3*/*pca4*.

Interestingly, also a further putative ferric reductase gene related to *pca1* is strongly up-regulated upon plant recognition (Triha1_76871; 21.2fold) as is a copper transporter gene closely related to *pca2* (Triha1_83588; 30.8fold). Both genes are not located in the genomic vicinity of the PCA cluster.

A SNP analysis of differences between *T. harzianum* B97 [48] and the publicly available sequence of *T. harzianum* (*sensu stricto*) CBS226.95 [41] revealed no intragenic, no intergenic and no nonsynonymous SNPs in the region of the PCA cluster. Only one synonymous SNP was detected in 513506 and one in 513501 in the 5’ UTR, hence strengthening the identification of *T. harzianum* B97 as *sensu stricto*. Consequently, we will refer to the *T. harzianum* CBS226.95 protein IDs and sequences hereafter.

In order to gain information on the potential function of the PCA cluster, we performed domain analysis and checked homologous genes in other fungi. PCA1 comprises a NADPH oxidase domain (cd06186) catalyzing the generation of reactive oxygen species (ROS) as well as a ferric reductase like transmembrane component (pfam01794) and is related to *A. fumigatus* FRE7, which is regulated by veA [56], upon response to Fe starvation [57] and during hypoxia [58]. PCA2 contains a copper transporter domain (pfam04145), which may be involved in oxidative stress protection or pigmentation and is related to putative low affinity copper transporters in *Aspergilli*. PCA3 is a member of the transferase superfamily (cl23789), which comprises enzymes that catalyze the first committed reaction of phytoalexin biosynthesis, but also trichothecene 3-0-acetyltransferase. PCA3 represents a homologue of the *Fusarium* trichothecene 3-O-acetyltransferase Tri101. This enzyme is known to have a function in self-protection of trichothecene producing fungi like *Fusarium pseudograminearum* [59]. Although trichothecene production was reported for *Trichoderma* [60,61], predominantly in the Brevicompactum clade [62] comprising *T. brevicompactum* or *T. arundinaceum*, but not for species of other clades, we checked for the presence of genes associated with trichothecene production. Besides PCA3/Tri101, the *T. harzianum* genome comprises homologues of the trichodiene oxygenase TRI4 (Triha1_99853), the transcription factor TRI6 (Triha1_514309, distant relationship), the transcription factor TRI15 (Triha1_1233) and the isotrichodermin C-15 hydroxylase TRI11 (Triha1_535230). None of these genes is regulated upon recognition of a soy plant in our experiment (supplementary file 1).

PCA4 contains an NAD/NADP octopine/nopaline dehydrogenase (pfam02317) domain as well as a glutamate synthase or related oxidoreductase domain (cl28234), which may be involved in amino acid transport and metabolism. PCA4 has no homologues in *Aspergilli*. PCA5 is a major facilitator superfamily transporter (pfam07690), related to a cycloheximide resistance protein. PCA6 belongs to the superfamily of S-adenosylmethionine-dependent methyltransferases, class I (cl17173). PCA7 has a Cytochrome P450 domain (cl12078), which may be involved in the degradation of environmental toxins. It is relatead to the *Fusarium* isotrichodermin C-15 hydroxylase Tri11, but is no direct homologue according to reciprocal best hit analysis. In contrast to *Trichoderma brevicompactum* [61], *T. harzianum* does not comprise a trichodermin cluster in its genome, as also outlined above. The homologue of PCA7 in *A. nidulans*, STCF, is a putative sterigmatocystin biosynthesis P450 monooxygenase with a predicted role in sterigmatocystin/aflatoxin biosynthesis.

Due to the tremendous up-regulation of the PCA cluster upon plant recognition, we were interested whether the genes of this cluster or their homologues in *T. virens* (which has the complete PCA cluster) are regulated in the presence of a plant as well. Therefore, we evaluated and/or re-analyzed available transcriptome data from earlier studies with maize or tomato, which however were carried out with different growth conditions [63–67]. In most cases, the *pca*-genes and their homologues in *T. virens* were not significantly regulated (>2fold, p-value<0.05) during plant colonization and were not detected in the secretome associated with plant contact. Only in one study [64], down-regulation of four of the seven PCA cluster genes was detected in a hydroponic interaction system in the colonization phase. (for details see supplementary data 1, supplementary file 2). Hence, we conclude that the PCA-cluster is particularly important during early plant recognition under conditions simulating solid phase interaction with the plant.

### The PCA cluster region comprises multiple occurrences of a novel DNA motif

Due to the striking up-regulation of the PCA cluster genes upon plant recognition, we were interested whether a common DNA motif might be present in the promotors of the genes. To this end, we analyzed the promotors of all *pca* genes for promotor motifs using MEME. Interestingly, we found a palindromic motif (Figure 3A,C) in the intergenic regions between *pca1* and *pca2* as well as *pca3* and *pca4*. This motif is present twice in each region and was not yet characterized in fungi.

### The PCA cluster is required for plant root colonization

We further asked whether the early and strong transcriptional response of the PCA cluster genes upon plant recognition would be predictive of a role in colonization, which is crucial for plant-fungus interaction and plant protection [26]. For construction of deletion strains we chose *pca1*, the putative NADPH oxidase, since previous work revealed an involvement of ROS – directly or indirectly – in plant-fungus interaction [32,33]. Additionally, we prepared a null mutant of *pca5*, encoding a transporter potentially involved in signal compound emission. Both mutant strains were viable and showed no striking growth defects (Figure 4A).

**Figure 4.**
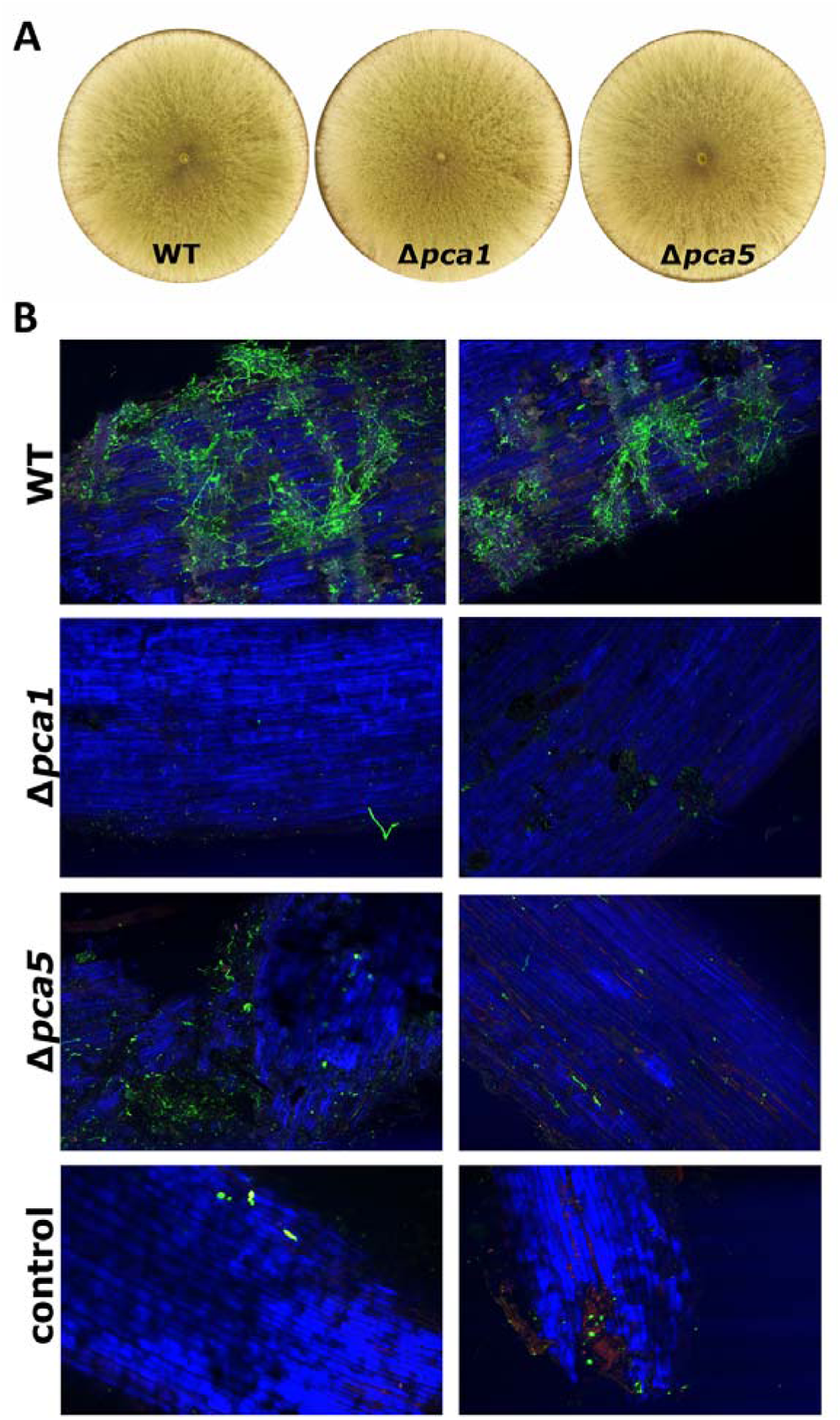
Colonization of soybean roots by *T. harzianum* B97 wildtype and mutant strains Δ*pca1* and Δ*pca5*. (A) Phenotype of recombinant strains (Δ*pca1* and Δ*pca5*, representative plates shown) grown for 72 hours in light on malt extract media. (B) Uninoculated roots were used as control. Fungal mycelia on the soybean roots were stained with WGA-Alexa Fluor488®. CLSM micrographs are showing B97 hyphae as green fluorescent colonizing the roots of wildtype but hardly detectable with both mutant strains. The two panels for each mutant (left panel: strains pca1_13Aa and pca5_28Aa; right panel pca1_4Ba and pca5_27Ba) show representative pictures of independent transformants, selected from at least three biological replicates each and several sites of the respective replicate.

We tested the ability to colonize plant roots by inoculating soybean seeds with the mutant strains or the wildtype and evaluated the presence of fungal mycelia on young roots after 8 days (Figure 4B). Confocal microscopy was performed from at least three replicate assays and multiple sites per root with two independent deletion strains per mutation. Roots grown from uninoculated seeds were used as controls. This analysis showed that while the wildtype *T. harzianum* B97 efficiently colonized the root surface of soy plants in addition to those of wheat as shown in Figure 1A. Neither Δ*pca1* nor Δ*pca5* were able to colonize the root grown from seeds inoculated with the mutant strains and these samples rather resembled the uninoculated control (Figure 4). We conclude that the requirement of these two genes of the PCA cluster is representative for the importance of this cluster and its upregulation upon plant recognition for efficient colonization of plants by *T. harzianum* B97.

### The PCA cluster is specific to Trichoderma and subject to balancing or positive selection

Because of the obvious importance of the PCA cluster for plant recognition and colonization, we were interested in its conservation and evolution in fungi. Since the genomic region of the PCA cluster in B97 did not comprise a notable number of SNPs in comparison with *T. harzianum* CBS226.95, we will refer to the region in the latter strain in our further analyses and descriptions.

After identifying the genomic area of the PCA cluster in *T. harzianum*, we performed a blastn analysis with all dikarya genome sequences available at JGI (2140 genomes). These genome sequences cover the group of Sordariomycetes (528 genomes) very well, including numerous strains of the genus *Trichoderma*, which would allow for association to the well-studied clades of the genus [68]. All other groups of fungi in JGI mycocosm outside dikarya [69] were tested as well, but did not yield homologous sequences. Surprisingly, the search results did not reflect the expected relationships according to the known phylogeny of ascomycetes. Moreover, the cluster was not present in many *Trichoderma* species. Neither *Trichoderma* spp. outside of the Harzianum clade nor the common ancestor of the genus *Trichoderma*, *Escovopsis weberi* [70] or closely related species such as *Fusarium* spp. had this cluster, as revealed by only partial coverage of the cluster sequence in the genomes. Rather it was scattered among some species in the Sordariomycetes. However, good coverage of the cluster area was detected for *Metarhizium* species as well as for *Pestaliopsis fici* (Xylariales [71]) and *Talaromyces islandicus* (Eurotiomycetes [72] (Figure 5A and supplementary data 2 in supplementary file 2). All genes present in these fungi had high homologies and very low E-values compared to the genes in T. harzianum CBS226.95 as revealed by blastp analysis (supplementary file 2, supplementary table S7). Using the respective nucleotide sequences covering the whole clusters for phylogenetic analysis confirmed the close relationship of the cluster sequences (Figure 5B). In case of *Metarhizium* spp. we found coverage of the cluster for *M. anisopliae, M. robertii* and *M. brunneum*, representing the generalist species of the genus [73], but not in intermediate or specialist species, which is also the case for other secondary metabolite clusters in specialist species of *Metarhizium* [73]. Interestingly, the PCA cluster was not detected previously in *Metarhizium* [74] and does not overlap with the well characterized secondary metabolite clusters responsible for production of destruxin, ferricrocin or other known toxins [73,74]. Furthermore, this finding suggested, that the PCA cluster was acquired by *T. harzianum* via horizontal gene transfer (HGT), likely from *Metarhizium* spp.

**Figure 5.**
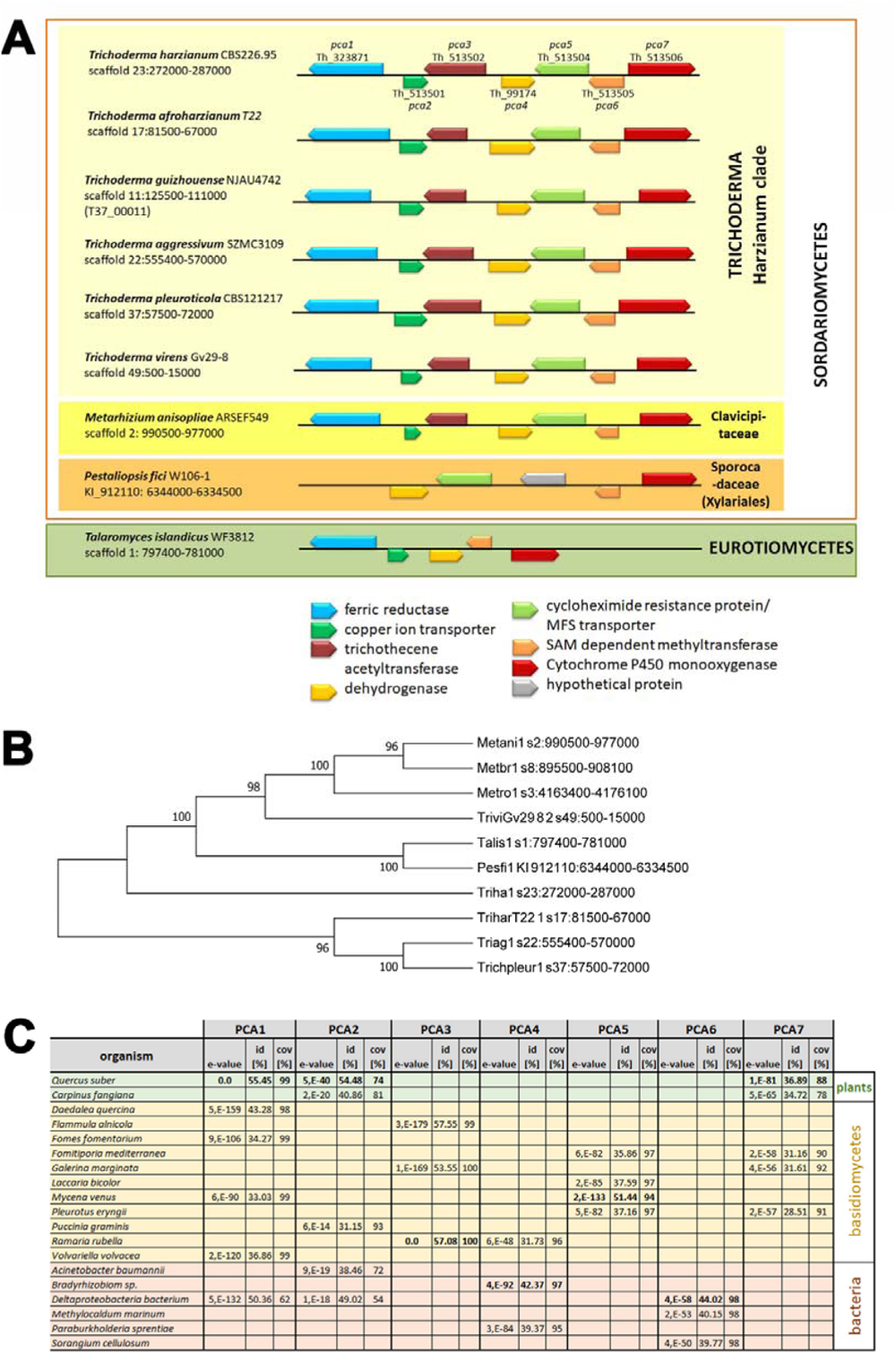
The PCA cluster, phylogenetic relationships and proteins related to its components. (A) Schematic representation of the clusters detected within selected representatives of *Trichoderma* spp. and outside the genus *Trichoderma*. For details on homology of the PCA-cluster homologues in these fungi, please see supplementary table S7 in supplementary file 2. (B) The evolutionary history was inferred by using the Maximum Likelihood method based on the Tamura-Nei model. The bootstrap consensus tree inferred from 1000 replicates is taken to represent the evolutionary history of the taxa analyzed. Branches corresponding to partitions reproduced in less than 50% bootstrap replicates are collapsed. The percentage of replicate trees in which the associated taxa clustered together in the bootstrap test (1000 replicates) are shown next to the branches. Initial tree(s) for the heuristic search were obtained automatically by applying Neighbor-Join and BioNJ algorithms to a matrix of pairwise distances estimated using the Maximum Composite Likelihood (MCL) approach, and then selecting the topology with superior log likelihood value. Sequences of *Metarhizium anisopliae* (Metani1), *Metarhizium brunneum* (Metbr1), *Metarhizium robertii* (Metro1), *Trichoderma virens* (TriviGV28_8_2), *Talaromyces islandicus* (Talis1), *Pestaliopsis fici* (Pestfi1), *Trichoderma harzianum* (Triha1), *Trichoderma afroharzianum* (TriharT22) and *Trichoderma aggressivum* (Triag1) originated from JGI mycocosm. (C) Blastp results of the respective protein sequences from *T. harzianum* B97 against the NCBI nr database with ascomycetes excluded. The top 100 hits for PCA1-7 were screened for most interesting similarities for this table.

Detailed analysis of the ocurrence of the PCA cluster in *Trichoderma* spp. genomes available in JGI Mycocosm revealed that it is only present in those species of the Harzianum clade of *Trichoderma*, but not in the evolutionarily younger clades (supplementary data 2 in supplementary file 2). Consequently, we investigated whether the PCA cluster might have been acquired by *T. harzianum* by HGT, which was obviously not the case. However, we found that the similarity of plant gene products (but not bacterial gene products) encoded in plant genomes like *Quercus suber* or *Carpinus fangipana* is due to their acquisition of PCA1, PCA2 and PCA7 homologues from ascomycetes by HGT. Analysis of Tajima’s D as a measure of evolution indicated balancing or positive selection for the proteins within the PCA cluster. For details on these analyses see supplementary material (supplementary data 2 and Figure S1 in supplementary file 2).

### Assessment of potential compounds associated with the PCA cluster

Phylogenetic analysis or blast searches did not reveal a cluster similar to the PCA cluster for which the associated compounds would have been determined. Antismash analysis of the genomic region spanning the PCA cluster did not yield a result, likely because the cluster does not comprise a “core” gene such as a terpene cyclase, PKS or NRPS. Since two genes in a genomic locus separate from the PCA cluster encode a PKS and an NRPS and both are somewhat up-regulated (see above), the cluster could be split with the core genes separated from the cluster. Antismash analysis of this separate locus indicated several possible products with 6-methylsalicylic acid reaching the highest similarity score. To evaluate the hypothesis of a split cluster, we performed CBlaster analysis [75] for identification of similar BGCs in other fungal genomes. In many cases, the species comprising the full or a partial cluster are pathogens like *Pyricularia* spp. or *Colletotrichum* spp. (supplementary file 3). While all 7 genes are present in part of *Trichoderma* spp. and in *Metarhizium*, many other species only comprise the genes encoding PCA4, PCA5 and PCA6, which hints at these proteins representing the core chemistry.

Comparison of the CBlaster BGC results using Clinker [76] was performed to elucidate whether PCA-like clusters in other fungi potentially comprise core genes like such encoding PKS or NRPS as well. As this was not the case (supplementary file 4), it is unlikely that the PCA cluster is split and that the locus comprising an NRPS and PKS encoding gene are associated with the PCA cluster. However, this also leads us to the conclusion that the chemistry responsible for the compound formed by the proteins encoded in the PCA cluster is somewhat cryptic and the alternative that a plant metabolite might be modified by these proteins could be considered.

## Discussion

Recognition of plants by fungi is crucial if the actual cry for help in the form of root exudates [77] is to be heard. Beneficial fungi of the genus *Trichoderma* positively impact plants at multiple levels, also using chemicals for achieving their effect [26], but most importantly, they trigger the systemic immune response of plants [18,78]. *T. harzianum* strains produce a number of secondary metabolites [79] with interesting bioactivity including antifungal activity [80], most of which are not yet assigned to specific biosynthetic gene clusters.

Although secondary metabolism and the gene clusters involved in biosynthesis of secondary metabolites are well studied in *Trichoderma* species [38,81], the plant recognition specific PCA gene cluster we found in this study was not described before. The enormous extent of the induction of this cluster, which exceeds the changes in transcript abundance of all other regulated genes, indicates that communication with the plant is associated with this induction, which we confirmed with investigation of two crucial genes of the cluster (*pca1* and *pca5*). Moreover, the fact that the presence of a plant elicits a response in terms of secondary metabolite production is in agreement with the finding that the secondary metabolite pattern of *T. harzianum* B97 is altered in the presence of a plant. Notably, the mutual influence of fungus and plant happens prior to physical contact and may hence involve volatile organic compounds (VOCs).

The striking impact of deletion of members of the PCA cluster on colonization of soybean roots strongly supports a function of the associated secondary metabolite(s) in plant-fungus communication. Such communication is vital for beneficial interkingdom-interactions, which was also shown for *Trichoderma* [26]. The beneficial effects of *T. harzianum* B97 on plant health are extensive, facilitating commercial application, and may involve an influence of secondary metabolites – including those of the PCA cluster – on innate immunity as shown previously for *Trichoderma* [30]. However, as root colonization is a prerequisite for efficient interaction, it can be concluded that the early stage recognition and communication represents the major function of the PCA cluster. This hypothesis is further strengthened by the very early induction of the cluster, within only 13 hours of proximity and without direct contact between plant and fungus.

Our analysis of gene regulation patterns specific for early plant recognition by *T. harzianum* B97 revealed a significant enrichment of genes involved in detoxification by export, which is likely achieved by transporters. Therefore, we investigated the role of PCA5 in plant interaction and indeed found that this transporter is crucial for colonization. This finding is in agreement with the hypothesis, that the chemical communication with the plant as initiated by induction of the PCA cluster is dependent on export of secondary metabolites serving as signaling molecules by PCA5.

The PCA cluster comprises three genes putatively involved in biosynthesis or modification of secondary metabolites. Among the genes regulated in response to the presence of the plant, we found several more putative permeases and transporters, but no gene which might encode a transcription factor. Hence, it remains to be demonstrated, how the coordinated induction of the PCA cluster genes is achieved.

Considering the NADPH oxidase domain of PCA1, a relevance of an NADPH oxidase involved in ROS production for plant-fungus interaction was previously shown for *T. atroviride* [82]. For *F. oxysporum*, NADPH oxidase was found to be essential for chemotropic response to the presence of a plant [33].

The compound formed by the PCA cluster proteins or with their contribution remains to be determined, but the function of part of the proteins allows for an estimation of potential outcomes. PCA4, as an opine dehydrogenase/synthase, may be involved in condensing an amino acid (often arginine) with an alpha-keta acid such as pyruvate and reduction of the formed imine to a secondary amine. Such compounds are involved in Crown-Gall formation after *Agrobacterium tumefaciens* infection in plants and to some extent opine-related molecules serve as signaling compounds [83]. Moreover, such compounds could potentially be methylated and oxygenated by PCA6 and PCA3.

Another possibility is that the fungus may use the compounds to obtain metals from the host – which is corroborated by the presence of the ferric/cupric reductase PCA1 and the copper transporter PCA2 in the cluster. The recently reported mechanism involving opine-type proteins was so far only described for bacteria [84]. In the bacterial pathogen *Staphyllococcus aureus* the compound staphylopine (related to opines and derived from pyruvate) is involved in metal acquisition [85] in a mechanism which includes specific recognition of the metal bound protein [86]. According to this hypothesis, an octopine-like compound would be made by PCA4, then methylated and acetylated and exported/imported to the host. Once the metal-bound molecule is re-imported, then the metal reductase would release the metal from the siderophore. While such a process would reflect a form of communication, the presence and (positive or negative) role of such a mechanism in inter-kingdom interaction remains to be clarified.

Currently we cannot anticipate for the biosynthesis of which compound the PCA cluster is responsible and if it may be harmful or toxic, because secondary metabolite clusters related to the PCA cluster were not characterized before. However, since the problematic *T. brevicompactum*, which produces harmful toxins [87], does not comprise the PCA cluster, it is unlikely that this cluster causes production of harmful chemicals during early plant recognition.

For plant associated fungi, like those of the genus *Trichoderma*, HGT seems to be a rather common phenomenon, especially concerning genes involved in production of secondary metabolites [88]. Hence, we tested the hypothesis that the discordance of the phylogenetic trees of the PCA cluster proteins with ascomycete phylogeny might reflect HGT of the PCA cluster. However, this was not the case – in contrast, this analysis suggested that plants had acquired part of the cluster from fungi.

Nevertheless, the similarity of the *T. harzianum* PCA cluster with that in *Metarhizium* spp.[89] suggests potential functionalities. *Metarhizium* spp. are known endophytes [90,91] and insect pathogens infecting hundreds of species [92,93]. In a tripartite interaction, *M. robertsii,* which comprises the PCA cluster, transfers nitrogen from insects they had infected to their plant hosts [94]. A comparable situation was shown for *Laccaria bicolor* which associates with pine and spruce and transfers nitrogen from collembola in soil to the roots it colonizes [95]. Recently, an impact of a strain belonging to *T. afroharzianum*, a species closely related to *T. harzianum*, on the interaction of a plant with pathogenic insects was shown [96,97]. Although this interaction was rather indirect via modulation of the gut microbiome, also numerous direct antagonistic effects on insects by *Trichoderma* are known [98]. Consequently, an ecological function of *Trichoderma* comparable to that of *Metarhizium* shown by Behie and colleagues [94] seems likely.

Since availability and uptake of different nitrogen sources considerably influences secondary metabolism [99], the upregulation of the *niiA* and *niaD* homologues upon plant recognition in *T. harzianum* B97 may not only reflect nitrogen transport to the plant, but could also be a sign for increased efforts for production of specific secondary metabolites for plant communication. Notably, although several genes associated with secondary metabolism are down-regulated, besides the strong induction of PCA cluster genes, also an NRPS encoding gene and a PKS encoding gene are slightly upregulated. These obvious shifts in secondary metabolism are in agreement with the altered pattern we observed due to the presence of the plant.

Species of the Harzianum complex are supposed to be the most common endophytic species in tropical trees [100], with speciation leading to habitat preferences of soil or endophytism [21]. Among the *Trichoderma* spp of the Harzianum clade, which comprise the PCA cluster, there are also endophytically growing ones like *T. endophyticum* and *T. afrasin* [21]. The upregulation of the *niiA* and *niaD* homologues, which are involved in nitrogen uptake upon recognition of the plant (*nit3* and *nit6*) indicates that nitrogen metabolism also may play a role in the interaction of *T. harzianum* with the plant, although potentially also other sources of nitrogen are used than by *Metarhizium*, which degrades killed insects to deliver nitrogen to the plant [94]. Consequently, the PCA cluster is likely to support beneficial communication to the plant at an early stage of colonization, which may involve pretending to be an arriving endophyte delivering additional soil/organic nitrogen. Since the PCA cluster is crucial for colonization by *T. harzianum*, presence of this cluster in the fungus is highly likely to contribute to the high efficiency of members of this clade in biocontrol applications. Accordingly, microbiome analysis revealed co-occurrence of *Trichoderma* and *Metarhizium* species in the rhizospere at high yielding field sites [101] or with banana plants [102].

In summary, we developed a strategy to simulate conditions of chemotropic response, which allowed us to detect a secondary metabolite cluster essential for communication with a plant enabling efficient colonization of the root surface (Figure 6). While revealing an intriguing new aspect of plant-fungus interaction, this finding can also be applied to evaluate the specificity of this regulation for prediction of high efficiency biocontrol capacity during strain screening. Thereby, the presence of the cluster as well as its high induction level represent promising features for diagnostic tests in strain screening programs. Additionally, future identification of the compound produced by the PCA cluster, which facilitates colonization, may support and enhance plant-fungus interaction of diverse biocontrol agents and enable plant-protection of plant varieties with otherwise insufficient response to *Trichoderma*-based biocontrol agents.

**Figure 6.**
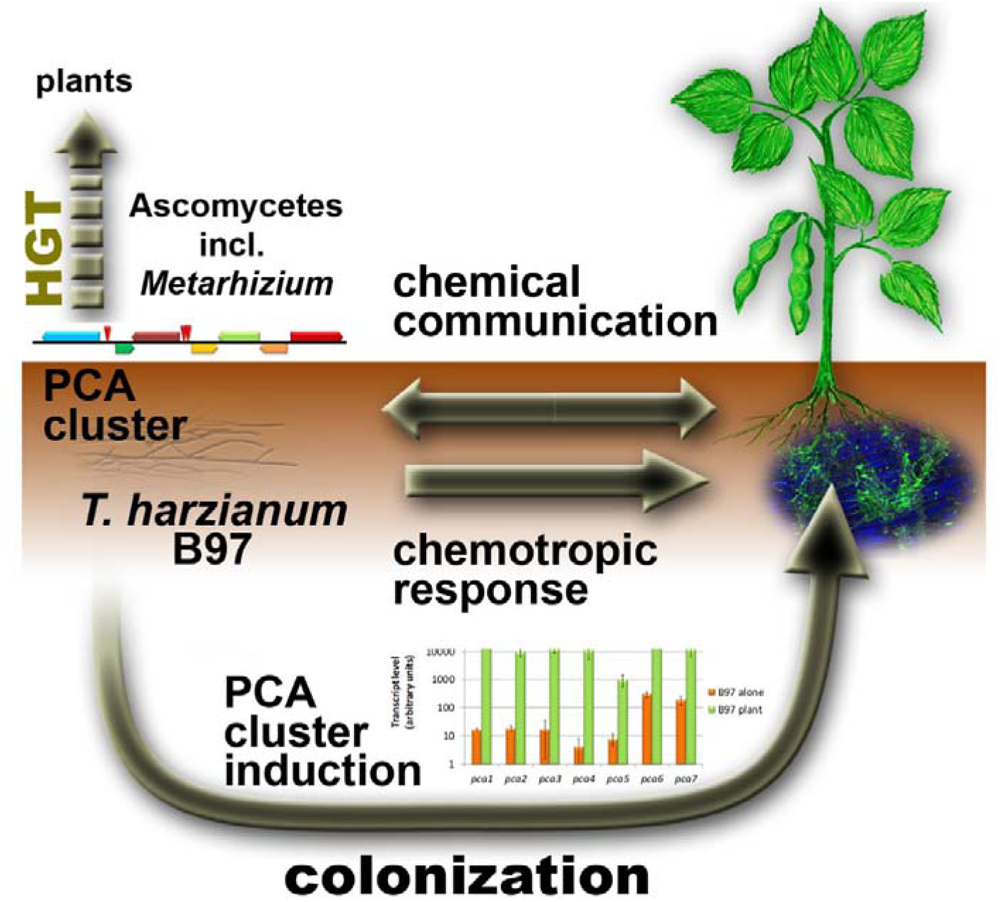
Schematic representation of the reaction of *T. harzianum* B97 to soy bean. *T. harzianum* chemotropically responds to the presence of a soy plant. Chemical communication occurs both ways due to alteration of the secondary metabolite pattern of both *T. harzianum* B97 and the soy plant. The PCA gene cluster, which highly similar to a cluster in *Metarhizium*, is strongly induced upon plant recognition and essential for effective colonization of plant roots and required for efficient colonization. Individual homologues of cluster genes were acquired by HGT by plants from ascomycetes.

## Materials and Methods

### Strains and cultivation conditions

*T. harzianum* B97 [48] was used throughout the study. For RNA analysis, the strain was revived from long term storage on malt extract agar (3 % w/v). Plates containing modified Mandels Andreotti minimal medium [103] with 0.1 % (w/v) glucose as carbon source were inoculated with 10 µl of spore solution (10^8^ spores/ml) at 28 °C in constant darkness for 34 hours. The modified Mandels-Andreotti medium was prepared as follows: The mineral salt solution contained 2.8 g/l (NH_4_)_2_SO_4_ (21.19 mM) (ROTH, Karlsruhe, Germany), 4.0 g/l KH_2_PO_4_ (29.39 mM) (Sigma-Aldrich, St. Louis, USA), 0.6 g/l MgSO_4_·7H_2_O (2.43 mM) (ROTH, Karlsruhe, Germany) and 0.8 g/l CaCl_2_·2H_2_O (5.44 mM) (Merck, Darmstadt, Germany). The trace element solution contained 0.250 g/l FeSO_4_·7H_2_O (0.899 mM) (Sigma-Aldrich, St. Louis, USA), 0.085 g/l MnSO_4_·H_2_O (0.503 mM) (Merck, Darmstadt, Germany), 0.070 g/l ZnSO_4_·7H_2_O (0.243 mM) (Riedel-de Haen, Seelze, Germany) and 0.143 g/l CoCl_2_·6H_2_O (0.603 mM) (Sigma-Aldrich, St. Louis, USA) and the pH was adjusted to 2.0 with concentrated sulfuric acid. The culture medium was prepared combining 500 ml mineral salt solution, 480 ml milliQ water, 20 ml trace element solution, 0.0025 % (w/v) peptone from casein (Merck, Darmstadt, Germany) to facilitate germination, 0.1 % (w/v) D-glucose (ROTH, Karlsruhe, Germany) as carbon source and 1.5 % (w/v) agar-agar (ROTH, Karlsruhe, Germany).

Plates were covered with cellophane in order to facilitate harvesting of mycelia. For recognition analysis, the roots of soy plants in the second leaf stage (19 days old) were washed 4 times with sterile distilled water and were placed on the plates with *T. harzianum* B97 in 3 cm distance from the growth front. After further incubation for 13 hours (corresponding to the time for recognition in the chemotropic assay) in darkness, fungal mycelia were harvested for RNA isolation and agar slices from the same area were excised for evaluation of secreted metabolite production. As controls, plates with fungus but no plant and plates with plant but no fungus were used. Five plates each were pooled per sample and three biological replicates were used.

### Construction of T. harzianum B97 deletion strains

Vectors for deletion of *pca1* and *pca5* were constructed by yeast recombination cloning using *hph* marker constructs with 1kb flanking regions as described previously [104]. Protoplast transformation was used for deletions in *T. harzianum* B97 parental strain with 10 mg/ml lysing enzymes (*Trichoderma harzianum*, Sigma # L-1412) and 150 µg/ml hygromycin B (Roth, Karlsruhe, Germany) for selection. Absence of the gene of interest was confirmed by PCR with primers binding inside the deleted region. In parallel, a PCR with standard primers amplifying *tef1* was performed to exclude a false negative result due to insufficient DNA quality or failed PCR assay. A list of primers used is shown in supplementary table S8 in supplementary file 2. The independent mutant strains pca1_4Ba, pca1_13Aa (lacking the *pca1* gene), pca5_27Ba and pca5_28Aa (lacking *pca5*) were used for further analyses and yielded consistent results.

### Surface sterilization of seeds and in-vitro culture of soybean plants

The soybeans (*Glycine max* (L) Merr., variety ES TENOR, Die Saat, Austria) were obtained from RWA Austria. For the surface sterilization, the soybeans were soaked in 70 % ethanol for 1 minute and then rinsed 3 times with sterile distilled water. Afterwards, the soybeans were transferred into a sterile beaker containing Danklorix (2.8 % sodium hypochlorite (w/w), Colgate-Palmolive, Vienna, Austria) and Tween 20 (Roth, Karlsruhe, Germany) and stirred for 3 minutes. The soybeans were then washed five times with sterile distilled water. The excess water was removed by placing soybeans on sterile paper tissue.

For the pre-germination, the surface sterilized soybeans were placed on sterile paper tissue soaked with tap water and incubated at 26 °C and 16 hours light:8 h darkness for 5 days. The seedlings were transferred to the *in-vitro* culture, which contained diluted Murashige & Skoog medium (to 0.5 concentration; Duchefa Biochemie, Haarlem, The Netherlands) and 0.8 % (w/v) Daishin Agar (Duchefa Biochemie, Haarlem, The Netherlands) at pH 5.8. The *in-vitro* cultures of soybean plants were further incubated at the same condition for two more weeks.

### Preparation of plant root exudates

After surface sterilization the soybeans were planted in sterilized perlite (premium perlite 2-6, Gramoflor GmbH, Germany). The soybeans were kept in a plant culture room at 26 °C with 16 hours light and 8 hours darkness. After approximately 3 days of emergence, the plantlets were then allowed to grow for further 2 weeks till the second leaf stage was achieved. The plantlets were recovered from the perlite carefully and washed gently under running water to remove the perlite. Afterwards, at least 300 plantlet roots were submerged in 500 ml sterile milliQ water and kept for 2 days at room temperature. The obtained root exudates were filter sterilized through Thermo Scientific Nalgene Syringe Filter with 0.2 µM pore size and stored at −80 °C.

### Isolation of total RNA

For isolation of total RNA, mycelium of the growth front from 5 replicate plates was pooled and frozen in liquid nitrogen. Three biological replicates were used with 5 pooled plates each. Samples were then treated as described previously [105] using the QIAGEN plant RNA kit (QIAGEN, Hilden, Germany). RNA quality and integrity were checked using Bioanalyzer 2100 (Agilent). Only high quality RNA was used for further analyses.

### Transcriptome analysis and bioinformatics

Sequencing of samples along with cDNA preparation was done in collaboration with VetCORE (Vienna, Austria) using Illumina HiSeq 50bp single-read sequencing. The software BWA [106] was used for mapping to the genome data of *T. harzianum* (JGI mycocosm; https://genome.jgi.doe.gov/Triha1/Triha1.home.html) [41]. Quality control of sequencing data was done using FASTQC and Trimmomatic [107] was used for trimming of reads. HTseq was applied for read counting [108]. The software samtools was used for data processing [109] and the limma package as implemented in R [110] was used for determination of statistically significant differential expression (>2fold, p-value threshold 0.01) along with the voom method [111], quantile normalization, linear model fitting (with lmFit) and empirical Bayes methods of assessing differential expression (with eBayes; [112]). Comparison of gene expression patterns between biological replicates yielded significance scores of ≥0.979 for both sample sets. Sequence data are available at NCBI GEO (Gene expression omnibus) under the accession number GSE229209.

HCE3.5 [113] was applied to perform hierarchical clustering with default settings and the Poisson correlation coefficient as the similarity/distance measure. FunCat (Functional category) analysis was done with the FungiFun2 online tool [114] based on bidirectional best hit analysis with *T. atroviride*.

### Analysis of chemotropic responses

The freshly grown spores from a 4 days old culture were recovered and dissolved in 1 ml spore-solution (0.8 % NaCl and 0.05 % Tween 80). After separation of mycelia by centrifugation through glass wool, the spore solution was centrifuged at 8000 g for 2 minutes, the supernatant was discarded, and the spore pellet was resuspended in 1 ml sterile milliQ water. For the chemotropism assay, the spore solution was adjusted to 10^8^ spores per ml with sterile milliQ water. Peptone from casein (Merck, Darmstadt, Germany) was used as germination stimulator in 0.5 % water agar. The concentration of peptone from casein was optimized to 0.0025 % (w/v). After 13 hours of incubation at 28°C in darkness, germling orientation was monitored for at least 400 germlings per sample and chemotropic index was calculated as described earlier [32].

### Analysis of patterns of secreted metabolites

Analysis for alteration of secondary metabolite patterns in the presence of a soy plant was essentially done as described previously [24,115]. Therefore the same conditions as applied for transcriptome analysis as outlined above were used (Figure 1C). Application of high performance thin layer chromatography (HPTLC) and data visualization was performed as described in [24] except that separation was done with chloroform and 1 mM trifluoroacetic acid in methanol.

### Analysis of colonization by T. harzianum B97 and recombinant strains

Seeds were surface sterilized in 70% ethanol for 7 minutes and rinsed for 3 minutes with sterile milliQ water. Seeds were then put onto MEX (malt extract) plates containing either *T. harzianum* B97, two B97Δ*pca1* and two B97Δ*pca5* strains or only MEX without fungus as negative control. Seeds were then placed in sterile magenta boxes containing soil mixture (1:1:1 perlite, sand, potting soil and 25 ml of sterilized tap water), which was autoclaved twice. After 8 days at 22°C under 12 hours light:12 hours darkness conditions, plants were harvested, and roots stained in 15ml phosphate buffered saline (PBS, pH 7.2) containing 5µg/ml wheat germ agglutinin (WGA)-AlexaFluor488 conjugate (Life Technologies, USA) and incubated for 2 hours at 37 °C before rinsing three times with PBS. Eight biological replicates were grown and treated for wild-type and controls, respectively. In case of mutant strains, each of the two independent mutants per gene-deletion was grown in 4 replicates, resulting in 8 replicate assays per mutation. Of those replicates, six were selected for microscopic analysis. On every replicate/root, several sites were monitored to ensure consistency of the results. All observations were carried out using a confocal microscope (Olympus Fluoview FV1000 with multi-line laser FV5-LAMAR-2 and HeNe(G)laser FV10-LAHEG230-2, Japan). Observations with the confocal microscope were done at objective lenses of 10x, 20x and 40x. Between 20 and 40 X, Y, Z pictures containing 20 to 60 scans were separately taken at 405, 488, 549 nm wavelengths in blue/green/orange-red channels respectively, with the same settings each time and normal light. The image analysis software Imaris was used on the confocal microscope to visualize 3D reconstructions. X, Y, Z pictures from different channels were then merged using the Image J software (version 1.47v), and Z project stacks were then used to create the pictures as described earlier [116].

### Phylogenetic analysis, clusters, HGT and evolution

For phylogenetic analysis, protein and nucleotide sequences were obtained from the NCBI nr database or the genome sequences available at JGI mycocosm. Sequences were aligned using Clustal X or MEGA7 [117,118] with default parameters. MEGA7 was used for phylogenetic analysis using standard parameters, the Maximum likelihood method and 1000 bootstrap cycles.

Horizontal gene transfer was analyzed using the python tool HGTphyloDetect [119]. Blastp was set to at least 1000 hits to cover close and distant relationships. Besides the conventional workflow, also the scripts for close and distant relationships implemented in HGTphyloDetect were tested in every case. Tajima’s D was calculated using DnaSP6 [120].

Potential biosynthetic genes and clusters were analyzed using antismash [121] as implemented in Galaxy (antismash version 6.1.1) [122]. Search for similar biosynthetic gene clusters (BGCs) was performed by CBlaster [75]. The CBlaster results for the BGCs were compared using Clinker [75].

## Author contributions

MiS performed gene deletion and confocal microscopy, GL performed chemotropism analysis and RNA isolation, WH performed secondary metabolite analysis, SC supervised confocal microscopy, DG supported colonization analysis, ADZ contributed to editing of the manuscript, JG and RC contributed to bioinformatic analyses and interpretation of the PCA cluster, MoS conceived the study, performed transcriptome analysis and wrote the final version of the manuscript. All authors read the manuscript and agreed to publication.

## Acknowledgements

We want to thank Stefan Böhmdorfer for providing access to the HPTLC analysis equipment and to the University of Natural Resources and Life Sciences (BOKU), Vienna for providing access to microscopy equipment.

Soybean samples were obtained from RWA Austria, Korneuburg, which is gratefully acknowledged. Work of MiS and MoS was supported by the Austrian Research Fund (FWF, grant P31464 to MoS), Work of WH was supported by the Lower Austrian Association for research promotion GFF (formerly NFB; grant LSC16-004 to MoS).

We want to acknowledge the JGI mycocosm for providing multiple fungal genome sequences for free public use, in part prior to publication.

## Conflicts of interest

The study was in part funded by Greencell, France.

## Availability of data

All data used for this study are available in the manuscript, its supplementary file and at the NCBI GEO online repository under accession number GSE229209.

## Supplementary material

**Supplementary file 1** contains data on differential gene regulation and statistics as well as gene annotations.

**Supplementary file 2** comprises replicate results for HPTLC analysis (supplementary figure S1), description of additional data on regulation of the PCA cluster genes (supplementary data 1 and supplementary tables S1-S4), analysis of horizontal gene transfer by *T. harzianum*, bacteria or plants along with phylogenetic analyses (supplementary data 2, figure S2 and supplementary tables S5 and S6). Moreover, this file contains supplementary table S7 (Blastp analysis of PCA cluster genes) and supplementary table S8 (Oligonucleotides used in the study).

**Supplementary file 3** contains the result of CBlaster analysis for the PCA cluster

**Supplementary file 4** contains the result of Clinker analysis for the PCA cluster

## Notes

https://www.ncbi.nlm.nih.gov/geo/query/acc.cgi?acc=GSE229209

### Summary of Updates

Additional analyses and supplementary materials provided, text revised for clarity, part of the figures revised and improved

