## supplementary file 2 for "Plant recognition by *Trichoderma harzianum* elicits upregulation of a novel secondary metabolite cluster required for colonization"

**Supplementary material**

**Table of contents**

**Supplementary figure S1** – Biological replicates of fungal plant interaction (HPTLC)	2

**Supplementary data 1	3**

*Regulation of the PCA-cluster genes and their homologues in* T. harzianum *and* T. virens	3

supplementary table S1A	3

supplemantary table S1B	3

supplementary table S2	4

supplementary table S3	4

supplementary table S4A	4

supplementary table S4B	5

**Supplementary data 2	6**

*Indications that individual genes of the PCA cluster presumably*

*originate from plants and bacteria*	6

Figure S2	7

*The PCA cluster was not acquired by horizontal gene transfer*	8

supplementary table S5	9

*The PCA cluster is largely subject to balancing evolution*	10

supplementary table S6	10

**Supplementary table S7** – Blastp analysis of PCA cluster genes	10

**Supplementary table S8** – Oligonucleotides used in the study	11

**References	12**

**Supplementary figure S1** – Biological replicates of fungal plant interaction (HPTLC)

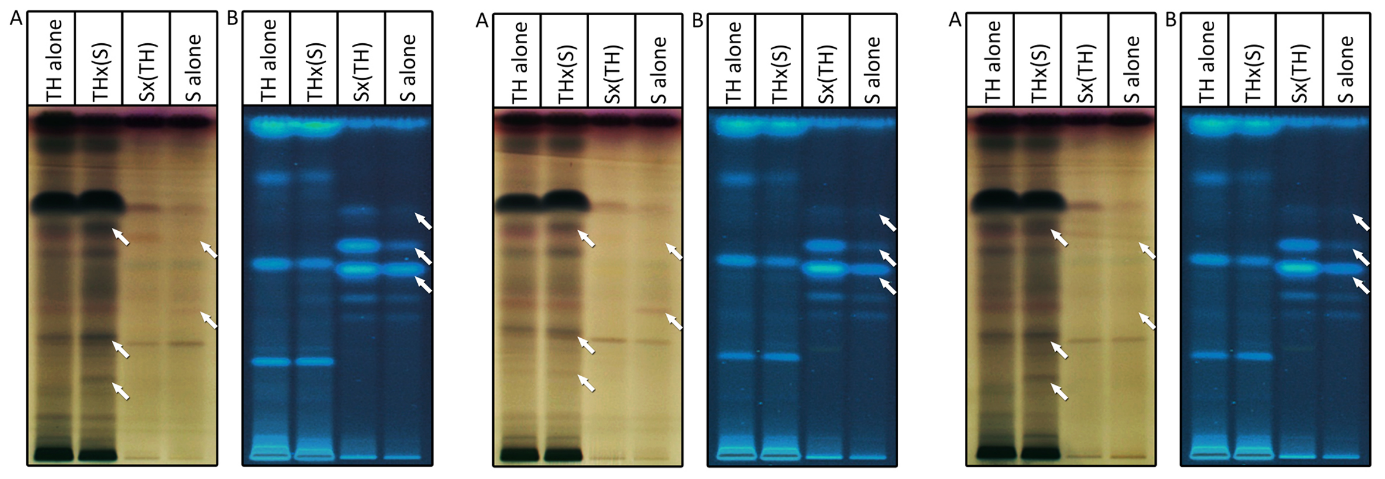

**REPLICATE 1	REPLICATE 2	REPLICATE 3**

**Supplementary figure S1**. **Assessment of *T. harzianum* B97 – plant interaction**. HPTLC analysis of *T. harzianum* B97 alone on the plate (TH alone), *T. harzianum* B97 in the presence of the root of soy plant (THx(S)), the root of soy plant in the presence of the fungus (Sx(TH)) and the root of soy plant alone (S alone). Two different visualizations are provided (A: Visible light after anisaldehyde derivatization and B: Remission at 366 nm) and show differentially secreted metabolites between interaction partners alone and in combination. Arrows mark bands with different abundance in the presence or absence of interaction.

Three different, independent biological replicates are shown.

**Supplementary data 1**

*Regulation of the PCA-cluster genes and their homologues in* T. virens

Considering the extreme induction of the PCA-cluster genes upon plant recognition, we were interested whether these genes are regulated during direct interaction with the plant as well. To get a broad picture on regulation, we re-analyzed different assays of plant interaction using GEO2R [1], which enables re-analysis and applies rigorous statistics on datasets submitted to NCBI GEO. For *T. virens*, which has the comlete PCA-cluster in its genome, we found in an RNAseq study that none of the pca genes was regulated significantly (p-value< 0.05, >2fold) upon interaction with tomato roots or maize roots (GSE64344; [2]) (supplementary tables S1A and S1B).

**Supplementary table S1A**. Transcript regulation of homologues of PCA-cluster genes in *T. virens* upon interaction with maize

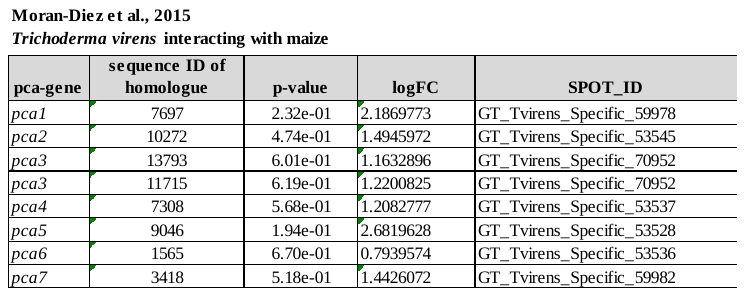

**Supplementary table S1B**. Transcript regulation of homologues of PCA-cluster genes in *T. virens* upon interaction with tomato.

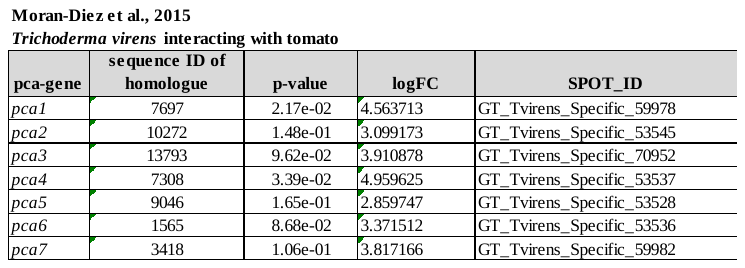

In a further RNAseq study [3], where a hydroponic system system was used, only *pca2*, *pca4*, *pca5* and *pca7* are represented. There, plant roots reaching into a nutrient solution were combined with pre-cultivated mycelia and shaken with 50 rpm. In contrast, our assay is based on solid media, in which diffusion is more important and volatile organic compounds (VOCs) may contribute to the interaction. The transcription levels of the detected genes remained below the threshold of 2fold significant regulation (p-value threshold <0.05) in the early so-called „recognition“ phase 6 hours after addition of mycelia to plant roots. Thirty hours after inoculation, all four genes are downregulated 7 – 20 fold compared to mycelium in the absence of plant roots (supplementary table S2).

**Supplementary table S2**. Transcript regulation of homologues of PCA-cluster genes in *T. virens* upon interaction with maize.

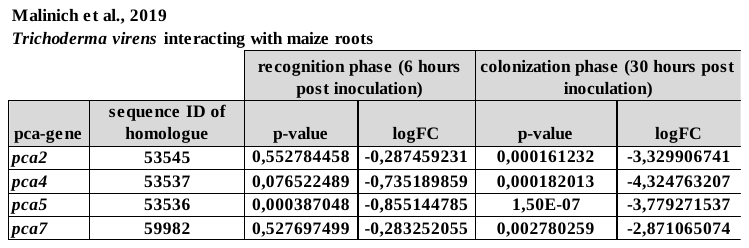

Earlier studies relied on the EST-dataset of the TrichoEST project, which comprised 12662 unique genes originating from 8 *Trichoderma* spp (including *T. harzianum*) cultivated under diverse conditions [4]. Screening of these sequences revealed that only *pca6* was represented among the sequences of *T. harzianum* and closely related sequences to *pca3* were present in *T. aggressivum* (supplementary table S3). This low representation supports the hypothesis that the PCA cluster is not relevant for axenic growth under diverse nutrient conditions and – limitations.

**Supplementary table S3**. Representation of PCA cluster genes among the TrichoEST clones

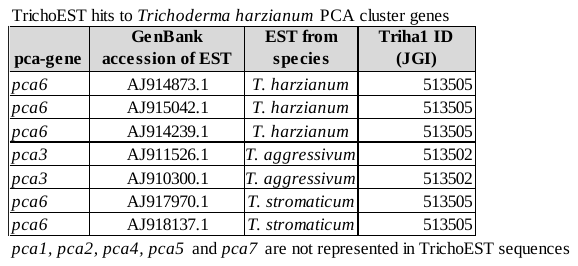

In a study based on these ESTs along with the whole genome of *T. virens*, interaction of *T. harzianum* and *T. virens* was analyzed for their transcriptomic response to tomato roots [5]. However, neither the *T. virens* sequences nor the *T. harzianum* sequences provided any indication of a regulation of the PCA-cluster genes during plant interaction (GSE29171; [5]) (supplementary table S4A, S4B).

**Supplementary table S4A**. Transcript regulation of homologues of PCA-cluster genes in *T. harzianum* upon interaction with tomato.

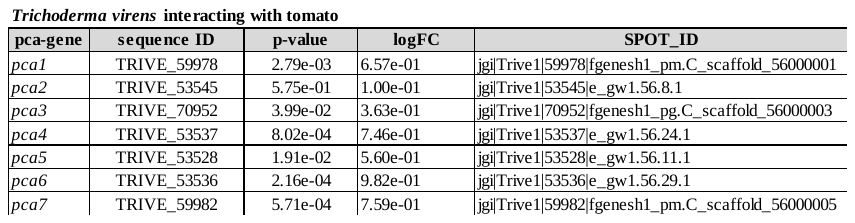

**Supplementary table S4B**. Transcript regulation of homologues of PCA-cluster genes in *T. virens* upon interaction with tomato.

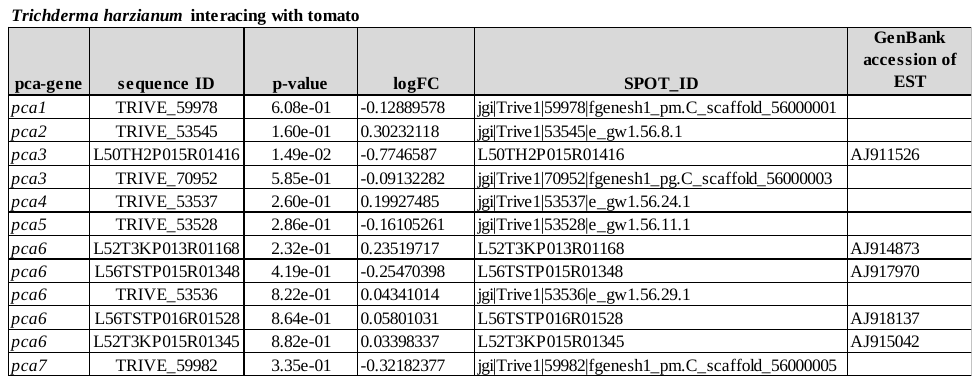

Finally, since PCA1, PCA2, PCA4 and PCA7 have signal peptides suggesting secretion, we screened the results of two studies on secreted proteins upon plant interaction of *T. virens* [6,7], but none of the gene products of the *T.virens* PCA cluster was represented in the respective secretome.

These data support the assumption that the PCA cluster is indeed specifically induced upon early plant recognition and only plays a minor role at best during later stages of fungus-plant interaction. However, given the diverging cultivation conditions applied in the screened studies, it cannot be excluded that under *in vivo* conditions in soil the relevance of the PCA cluster extends to the colonization phase and may be specific to a fungal species and/or to a plant host.

**Supplementary data 2**

*Indications that individual genes of the PCA cluster presumably originate from plants and bacteria*

Our detailed investigation of genome sequences confirmed that the full PCA cluster is indeed only present in the Harzianum clade [8] of *Trichoderma*, but not in the more ancestral Trichoderma clade or in the evolutionarily younger Longibrachiatum clade. These findings suggest that the PCA cluster was gathered from other organisms by HGT, likely after the split of the Harzianum clade from the Longibrachiatum clade in *Trichoderma*.

We screened the identified genomic locations for the presence of the seven genes of the PCA cluster and found all of them in the same order and orientation in the Harzianum clade as well as in *Metarhizium* spp., which supports the hypothesis of HGT. In *P. fici* and *T. islandicus* the cluster is not complete, with *pca1* and *pca2* or *pca3* and *pca5* missing, respectively (Figure 5A). Individual genes of the cluster did show hits with blast searches in *Trichoderma* spp. outside of the Harzianum clade as well as in several other species, but they were not assembled in clusters in the respective genomes.

We therefore asked whether the *pca* genes might be present in the genomes of other *Trichoderma* clades, yet not assembled in clusters. We performed phylogenetic analysis using at least three blast hits (if present and e-value <e-10; otherwise all available) for the predicted protein sequences encoded by each *pca* gene in various organisms. This analysis allowed us to delineate, whether in these species the cluster simply lost several components during evolution or if those blast hits merely represent unrelated genes with similar function. We used representatives of the Harzianum species complex [9] (*T. harzianum, T. guizhouense, T. afroharzianum*) and the Harzianum clade (*T. pleuroticola, T. aggressivum* and *T. virens*) [9]. From the further clades we picked representatives from the Longibrachiatum clade (*T. reesei*), the Helicum clade (*T. helicum*) [10], the Brevicompactum clade (*T. brevicompactum*) [11] and the Trichoderma clade (*T. atroviride*) [12] as well as from *M. anisopliae,* *P. fici* and *T. islandicus* and other fungi having potential homologues in their genomes. Moreover, we added closely related sequences revealed as potential homologues from plants and bacteria as detected in blastp analysis using NCBI Blastp and excluding ascomycetes.

Interestingly, the latter blastp search on the NCBIdatabase yielded also highly related sequences to bacteria and plants for individual PCA proteins (Figure 5B), with *Quercus suber* (cork oak, plant) homologues as best hit for PCA1, PCA2 and PCA7, basidiomycete proteins for PCA3 and PCA5 as well as bacterial proteins for PCA4 and PCA6, which again suggested that the components of the cluster were assembled from different donor species.

This first exploratory phylogenetic analysis revealed the PCA proteins presumably connected by evolution and enabled us to distinguish those which clustered with proteins only similar, but not homologous to the PCA proteins and hence unrelated to the cluster. Indeed, the PCA homologues from species comprising the cluster formed a separate clade from those with similarity to the PCA sequences, but not organized in a cluster. In a second step we then used the closest hit of respective protein sequences from this sister clade for repeated phylogenetic analyses (Figure S1 A-G).

This analysis confirmed that only the *pca* genes from species having the cluster form a clade together, while *pca*-related genes scattered over the genome formed separate clades. Interestingly, *T. atroviride* seems to have homologues of PCA1 and PCA3 (Figure S1A, C), since the respective proteins cluster closely with the genes assembled in clusters in *T. harzianum* and not outside this clade. Additionally, we also found that hits from fungi outside ascomycetes, as well as those from plants or bacteria clustered among the fungal sequences or as sister clades hence confirming the blastp results. Consequently, acquisition of individual genes even from different kingdoms seems possible, although the highly conserved structure of the cluster in *Metarhizium*, *Trichoderma* and in part *P. fici* and *T. islandicus* suggests assembly of the cluster in one of these genera.

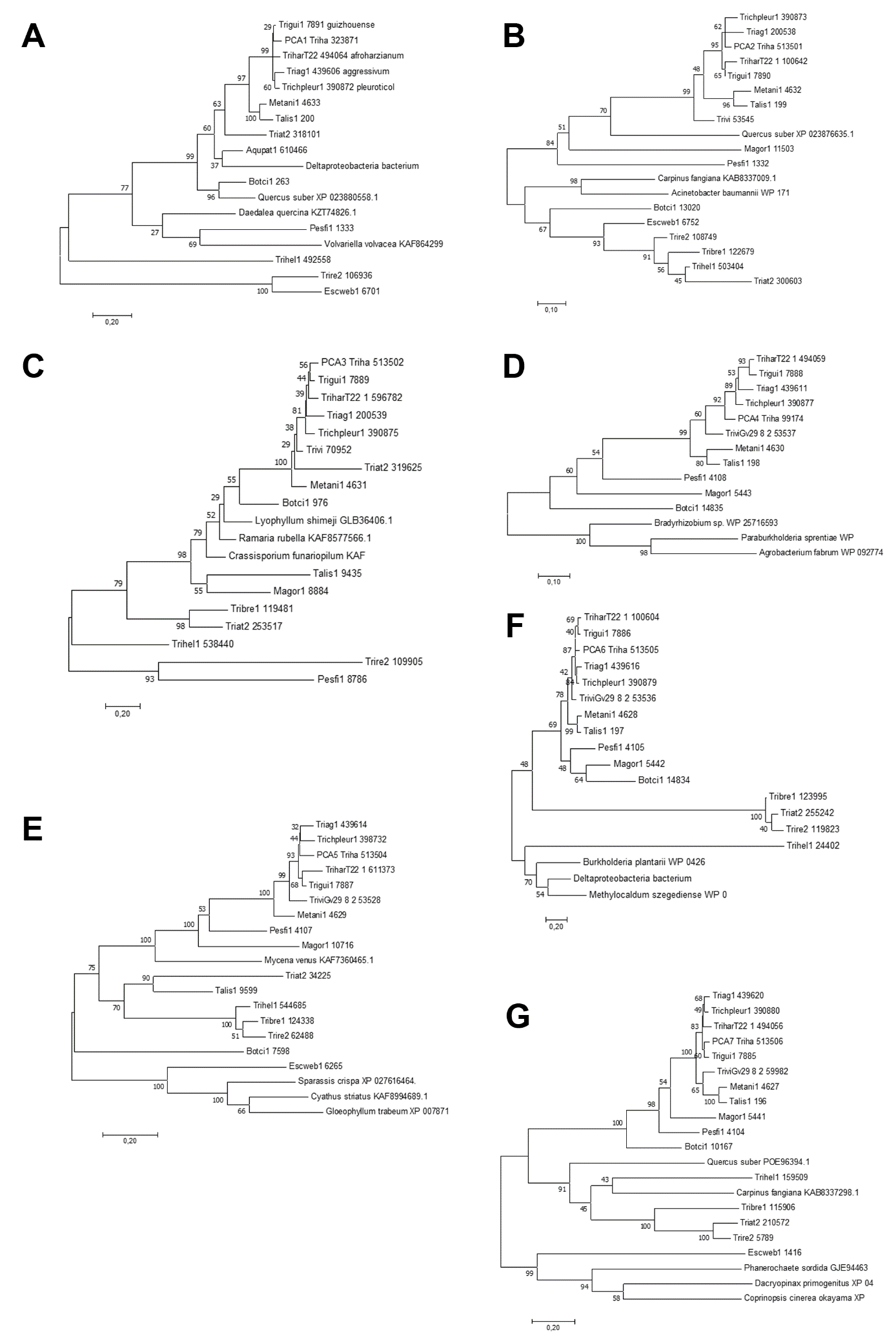

Figure S2. **Molecular phylogenetic analysis of the PCA cluster genes by the Maximum Likelihood method.** Maximum likelihood trees of homologues of (A) PCA1, (B) PCA2, (C) PCA3, (D) PCA4, (E) PCA5, (F) PCA6 and (G) PCA7 are shown. The evolutionary history was inferred by using the Maximum Likelihood method based on the JTT matrix-based model. The tree with the highest log likelihood is shown. The percentage of trees in which the associated taxa clustered together is shown next to the branches. Initial tree(s) for the heuristic search were obtained automatically by applying Neighbor-Join and BioNJ algorithms to a matrix of pairwise distances estimated using a JTT model, and then selecting the topology with superior log likelihood value. The trees are drawn to scale, with branch lengths measured in the number of substitutions per site. All positions with less than 95% site coverage were eliminated. That is, fewer than 5% alignment gaps, missing data, and ambiguous bases were allowed at any position.

*The PCA cluster was not acquired by horizontal gene transfer*

Phylogenetic analysis of predicted proteins within the PCA cluster as described above revealed some discordance with species phylogeny. Additionally, the unusual structure of the cluster, missing the typical biosynthetic polyketide synthases (PKS) or non-ribosomal peptide synthases (NRPS) demanded a closer look. Besides the PCA cluster, also a locus on a different scaffold comprising genes encoding an NRPS and a PKS could potentially contribute to biosynthesis of the metabolite associated with the PCA cluster. Consequently, we were interested whether this inconsistency is the result of horizontal gene transfer of part or all of the genes of the PCA cluster and whether the additional locus is concerned as well.

Using the package HGTphyloDetect [13], we investigated all protein sequences of the PCA cluster as well as the NRPS and PKS, which are regulated as well but located in a different genomic site. An alien index (AI) [13] of ≥ 45 was found to be a good indicator of foreign origin of a sequence [14]. For HGT among closely related organisms (e.g. eukaryote to eukaryote) the HGT index [13] was applied in order to capture potential HGT events for example from *Metarhizium* to *Trichoderma* as well.

We found no support for the hypothesis that either the cluster or the PKS- or NRPS-encoding gene in the second locus were acquired by HGT (supplementary table S1). Nevertheless, the finding of obviously closely related proteins in other fungi, plants or even bacteria (Figure 5C) was still puzzling. Therefore we performed the same analysis with the respective sequences from plants and bacteria. No HGT could be confirmed for sequences related to *pca3*, *pca5* and those from bacteria or other fungi.

However, we found that *pca1*, *pca2* and *pca7* were indeed subject to horizontal gene transfer from fungi (*Botrytis* spp. and related species) to *Quercus suber* (supplementary table S1). In order to rule out that this result is merely caused by sequencing artifacts [15] in the published *Q. suber* sequence, we also tested the next best hits in plants for *pca7*. HGT was also detected for the *pca7* homologues in *Carpinus fangiana* and *Adiantum nelumboides* (supplementary table S1), hence rejecting the hypthesis of an artifact.

Interestingly, all individual genes of the PCA cluster have homologues in *Botrytis cinerea* (as evaluated using the JGI mycocosm genome databases for this species). However, the respective genes are not assembled in a cluster. Only *pca1* and *pca2* are next to each other in the genome. In *Q. suber* the homologues of *pca1*, *pca2* and *pca7* for which HGT is supported are not located in genomic vicinity, but on different scaffolds.

**Supplementary table S5**. Analysis of horizontal gene transfer using HGTphyloDetect.

|  | **species** | **gene** | **related to** | **alien-index** | **e-value** | **HGT** | **predicted Donor** |
| --- | --- | --- | --- | --- | --- | --- | --- |
| **PCA cluster** | *T. harzianum* | *pca1* |  | n.a. | - | no | - |
|  | *T. harzianum* | *pca2* |  | n.a. | - | no | - |
|  | *T. harzianum* | *pca3* |  | -55.91 | - | no | - |
|  | *T. harzianum* | *pca4* |  | 34.67 | - | no | - |
|  | *T. harzianum* | *pca5* |  | n.a. | - | no | - |
|  | *T. harzianum* | *pca6* |  | n.a. | - | no | - |
|  | *T. harzianum* | *pca7* |  | n.a. | - | no | - |
| **PKS** | *T. harzianum* | XP_024768017 |  | n.a. | - | no | - |
| **NRPS** | *T. harzianum* | XP_024768014 |  | n.a. | - | no | - |
| **distant homolo-gues of PCA cluster genes** | ***Quercus suber*** | **XP_023880558.1** | ***pca1*** | **460.52** | **0.0** | **yes** | ***Botrytis sp.*** |
|  | ***Carpinus fangiana*** | **KAB8337298.1** | ***pca1*** | **342.91** | **1.19e-149** | **yes** | ***Coleo-phoma sp.*** |
|  | ***Adiantum nelumboides*** | **MCO5585497.1** | ***pca1*** | **460.52** | **0.0** | **yes** | ***Meira spp.*** |
|  | *Deltaproteo-bacteria bacterium* | MAD86624.1 | *pca1* | n.a. | - | no | *-* |
|  | *Daedalea quercina* | KZT74826.1 | *pca1* | -460.52 | - | no | *-* |
|  | ***Quercus suber*** | **XP_023876635.1** | ***pca2*** | **160.72** | **1.59e-70** | **yes** | ***Botrytis sp.*** |
|  | *Ramaria rubella* | KAF8577566.1 | *pca3* | 0.00 | - | no | *-* |
|  | *Mycena venus* | KAF7360465.1 | *pca5* | -460.52 | - | no | *-* |
|  | ***Quercus suber*** | **POE96394.1** | ***pca7*** | **460.52** | **0.0** | **yes** | ***Zasmidium sp.*** |

n.a. not available i.e. calculation of alien index not successful, no support for HGT

*The PCA cluster is largely subject to balancing evolution*

The results of our phylogenetic analysis together with HGT analysis suggested that PCA cluster genes may have been lost during evolution in many fungal species, including *Trichoderma* spp. outside the Harzianum cluster. Therefore we were interested, whether evolutionary pressure may have caused this effect. To distinguish between random (neutral) evolution and non-random evolution like directional selection or balancing selection, we applied the Tajima’s D test [16]. In case of randomly involving sequences, no effect on the fitness or survival of an organism is expected and the frequenies of such neutral mutations would fluctuate through genetic drift. Mutations affecting the fitness or survival of an organism – such as for example lethal mutations – are non-neutral and under selection.

Calculation of Tajimas D from individual conding regions did not yield statistically significant results for synonymous sites, non-synonymous sites or overall except for synonymous sites of *pca5*. The Tajimas D (non-syn/syn) ratio shows tendencies towards balancing selection or a balance between mutation and genetic drift, for *pca6* and *pca7*, the negative values support a hypothesis of positive selection (supplementary table S2).

Hence we consider it likely, that the PCA cluster has a neutral or positive effect in species where it was retained, potentially in combination with the respective ecological niche, like for *T. harzianum*. Such a positive effect would also be in agreement with the acquisition of *pca7* by plants from fungi, although the benefit of this gene for plants remains to be determined.

Table S6. Tajimas D for the genes of the PCA cluster

| **gene** | **Tajimas D (nonsyn/syn) ratio** | **selection** |
| --- | --- | --- |
| *pca1* | -0,5636 | balance between mutation and genetic drift |
| *pca2* | 21,22982 | balancing selection |
| *pca3* | 25,01958 | balancing selection |
| *pca4* | 0,22122 | balance between mutation and genetic drift |
| ***pca5*** | **0,03811** | **balance between mutation and genetic drift** |
| *pca6* | -4,45727 | positive selection |
| *pca7* | -13,21566 | positive selection |

**Supplementary table S7** – Blastp analysis of PCA cluster genes in selected genomes (as shown in Figure 5A)

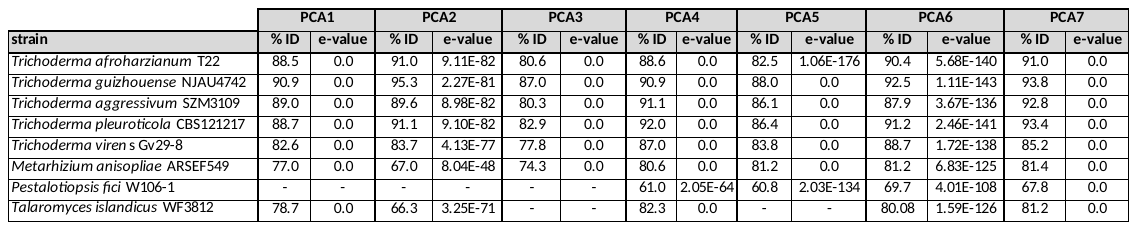

**Supplementary table S7**. Oligonucleotides used in this study

| **Name** | **Function** | **Sequence 5' - 3'** | **Protein ID** | **targtet gene** |
| --- | --- | --- | --- | --- |
| Bpca1del5F | creation of 5' flank of deletion cassette | 5’GTAACGCCAGGGTTTTCCCAGTCACGACGATGGTGGTGATTGTTGTG 3‘ | 323871 | *pca1* |
| Bpca5del5R | creation of 5' flank of deletion cassette | 5’ATCCACTTAACGTTACTGAAATCTCCAACGGTAAATGCGTTTCAAAG 3‘ | 323871 | *pca1* |
| Bpca1del3F | creation of 3' flank of deletion cassette | 5’CTCCTTCAATATCATCTTCTGTCTCCGACATTAAATGATACACAGGCTG 3‘ | 323871 | *pca1* |
| Bpca1del3R | creation of 3' flank of deletion cassette | 5’GCGGATAACAATTTCACACAGGAAACAGCCATTGTCATCTGCAGTAGAC 3‘ | 323871 | *pca1* |
| Bpca1screF | confirmation of deletion | 5’GGATGGACCTTACCCTTTATCG 3‘ | 323871 | *pca1* |
| Bpca1screR | confirmation of deletion | 5’ACCACAAACGAGTGCTGAAATC 3‘ | 323871 | *pca1* |
| Bpca5del5F | creation of 5' flank of deletion cassette | 5’GTAACGCCAGGGTTTTCCCAGTCACGACGTTGTCCGTTGTCCTATGGC 3‘ | 513504 | *pca5* |
| Bpca5del5R | creation of 5' flank of deletion cassette | 5’ATCCACTTAACGTTACTGAAATCTCCAACATCGCTTATTCGTTCGCAG 3‘ | 513504 | *pca5* |
| Bpca5del3F | creation of 3' flank of deletion cassette | 5’CTCCTTCAATATCATCTTCTGTCTCCGACGGTCCATTTGATAATAGAGAAG 3‘ | 513504 | *pca5* |
| Bpca5del3R | creation of 3' flank of deletion cassette | 5’GCGGATAACAATTTCACACAGGAAACAGCCTATTCACACCCAGAGCAC 3‘ | 513504 | *pca5* |
| Bpca5screF | confirmation of deletion | 5’CCGACGCAGGAAAGAAAC 3‘ | 513504 | *pca5* |
| Bpca5screR | confirmation of deletion | 5’ACAGATGTAGACGCAGCTGG 3‘ | 513504 | *pca5* |
