## Supplementary figures and images for "Plant recognition by *Trichoderma harzianum* elicits upregulation of a novel secondary metabolite cluster required for colonization"

### supplementary file 3

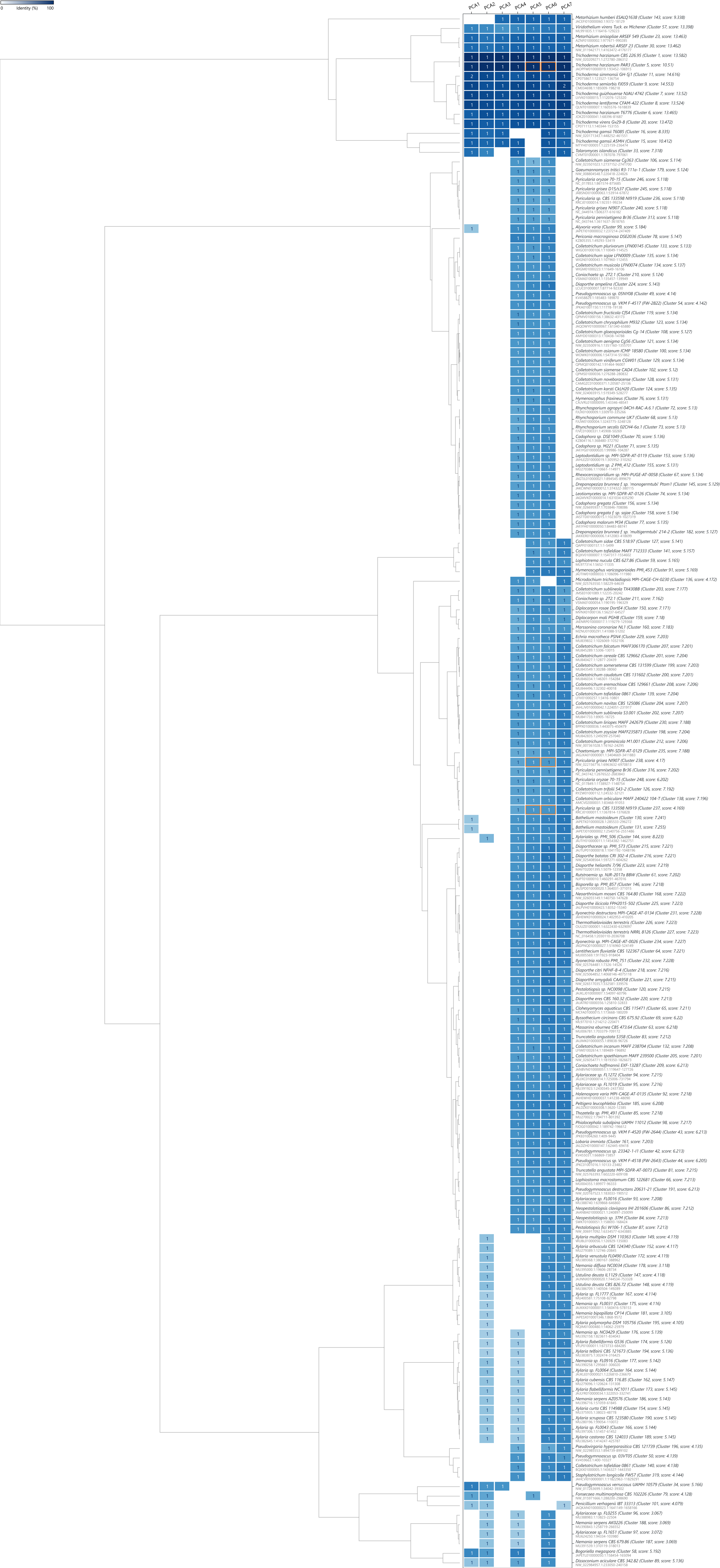
